# Inducible formation of fusion transcripts upregulates haploinsufficient CHD2 gene expression

**DOI:** 10.1101/2025.05.28.656657

**Authors:** Caroline Jane Ross, Yael Sarusi, Yoav Lubelsky, Rotem Ben-Tov Perry, Anat Mavashov, Shaked Turk, Moran Rubinstein, Igor Ulitsky

## Abstract

Modes of action of long noncoding RNAs (lncRNAs) are poorly understood. CHASERR is a broadly expressed lncRNA located immediately upstream of the promoter of the CHD2 gene. We show that antisense oligonucleotides (ASOs) targeting conserved motifs in CHASERR’s last exon induce the formation of a fusion transcript joining CHASERR to CHD2. This fusion transcript is exported to the cytoplasm and translated into full-length CHD2 protein. Deleting the same motifs in mice mimics the ASO effect, increasing CHD2 protein without causing the deleterious effects associated with full CHASERR ablation. The fusion transcripts are also expressed endogenously, induced in activated neurons, and their constitutive induction affects neuronal gene expression and chromatin accessibility. Perinatal introduction of the ASO into *Chd2*^+/–^ mice up-regulates CHD2 expression and alleviates behavioral phenotypes caused by CHD2 haploinsufficiency, providing a therapeutic route to CHD2 haploinsufficiency. This concept of targeting upstream genes with ASOs to induce transcript fusion can be extended to other gene pairs, and is thus a broadly relevant approach for increasing haploinsufficient gene expression.

## Introduction

High-throughput transcriptomic studies over the past decade have uncovered pervasive transcription of the mammalian genomes, which are now annotated to have a comparable number of loci producing long noncoding RNAs (lncRNAs) and protein-coding genes^1^. A growing number of lncRNAs have been associated with a wide variety of biological processes^2^. As a result, lncRNAs are increasingly recognized as therapeutically relevant entities. They can be harnessed as either full-length lncRNA molecules or functional fragments^3^, and can be also targeted by antisense oligonucleotides (ASOs)^4^ or genome-editing approaches^5^. The ASO-based approach already has promising results in pre-clinical studies and ongoing clinical trials^6^. However, for the majority of lncRNAs, the functions, if any, remain unclear. More challenging is the question of how they function and how their targeting leads to specific molecular and organismal outcomes, an understanding that remains limited even for lncRNAs with well-established functional importance^2^.

We have previously characterized *Chaserr*, a long noncoding RNA relatively conserved in sequence and genomic position, and present, to our knowledge, in all vertebrate species^7^. The high evolutionary conservation of *Chaserr* is likely due to its functional importance, as *Chaserr^-/–^* mice die shortly after birth, and *Chaserr^+/–^* mice exhibit various pleiotropic phenotypes across multiple tissues, often resulting in decreased survival. The functional importance of *CHASERR* was recently confirmed in humans as three *CHASERR^+/–^* probands were identified and shown to exhibit a syndromic early onset neurodevelopmental disorder that includes severe encephalopathy, shared facial dysmorphisms, cortical atrophy, and cerebral hypomyelination^8^. In humans and mice, loss of the *Chaserr* promoter (∼1 kb in *Chaserr* loss-of-function mouse model that we generated^7^ and ∼20 kb in the human cases^9^) leads to up-regulation of *CHD2*, a gene located on the same strand as *Chaserr* and ∼2 kb downstream of it. Deletion of different parts of the *Chaserr* gene or knockdown of the nascent transcript all lead to a >60% increase in *Chd2* mRNA levels, and at least as strong, and occasionally stronger, increase in CHD2 protein levels, in mouse and human cells^7,8,10^. In the case of *Chaserr* promoter deletion, where *Chaserr* transcription is lost throughout the locus, CHD2 up-regulation may be partially attributed to competition for a shared set of enhancers found in the large upstream gene desert. These enhancers form spatial contacts with both *Chaserr* and *Chd2* promoters, which are separated by ∼20 kb^7^. However, promoter competition can not fully explain CHD2 up-regulation, as it is also triggered by other *Chaserr* perturbations that leave the *Chaserr* promoter intact. These include deletion of the *Chaserr* gene body in mouse embryonic stem cells or knockdown using GapmeRs – antisense oligonucleotides that cleave their target transcripts. Therefore, transcription and/or RNA product of *CHASERR* are required for CHD2 regulation. The role of the RNA in this process is consistent with the unusually high evolutionary conservation of the *CHASERR* locus sequence compared to other lncRNAs.

*CHASERR* has a typical five-exon isoform in vertebrates, and much of its ∼1.1kb sequence is conserved during evolution^11^ (**Fig. 1A**). However, direct comparisons between distant *Chaserr* orthologs in human and zebrafish yielded no significant pairwise alignments. This motivated us to develop lncLOOM, an algorithmic approach for detecting short 6–15 nt sequences with conserved presence and order in a series of homologous sequences^10^. Application of lncLOOM to *Chaserr* sequences from 16 species identified a series of conserved elements in the beginning of the last exon of *CHASERR*. We refer to this region as the CHASERR Motif Containing Region (“MCR”), which includes two prominent RRAUGG (R=A/G) motifs that are clustered in multiple copies in the last exon of *CHASERR* throughout vertebrates^10^. We previously showed that blocking two of these elements with fully 2’MOE-modified ASOs increased CHD2 levels in Neuro2a cells^10^. In this study, we focus on the mode of action of these ASOs and the functional importance of the sequences that they target. We found that ASOs targeting the *CHASERR* MCR strongly induce the formation of a *CHASERR*-*CHD2* fusion transcript, which is naturally expressed at low levels, yet dynamically regulated during brain development and neuronal activation. The fusion contains the full coding sequence of CHD2 and can be translated to produce a full-length CHD2 protein. This fusion transcript is also up- regulated when the *Chaserr* MCR is deleted in a mouse model. Neurons from these mice have aberrant neuronal gene expression and chromatin accessibility, in particular in an activated state. Blocking the *CHASERR* MCR in a model of CHD2 haploinsufficiency up-regulates CHD2 protein and rescues phenotypes associated with Chd2 haploinsufficiency. Lastly, we show that the concept of ASO-mediated transcript fusion formation can be expanded to other tandem gene pairs.

**Figure 1.**
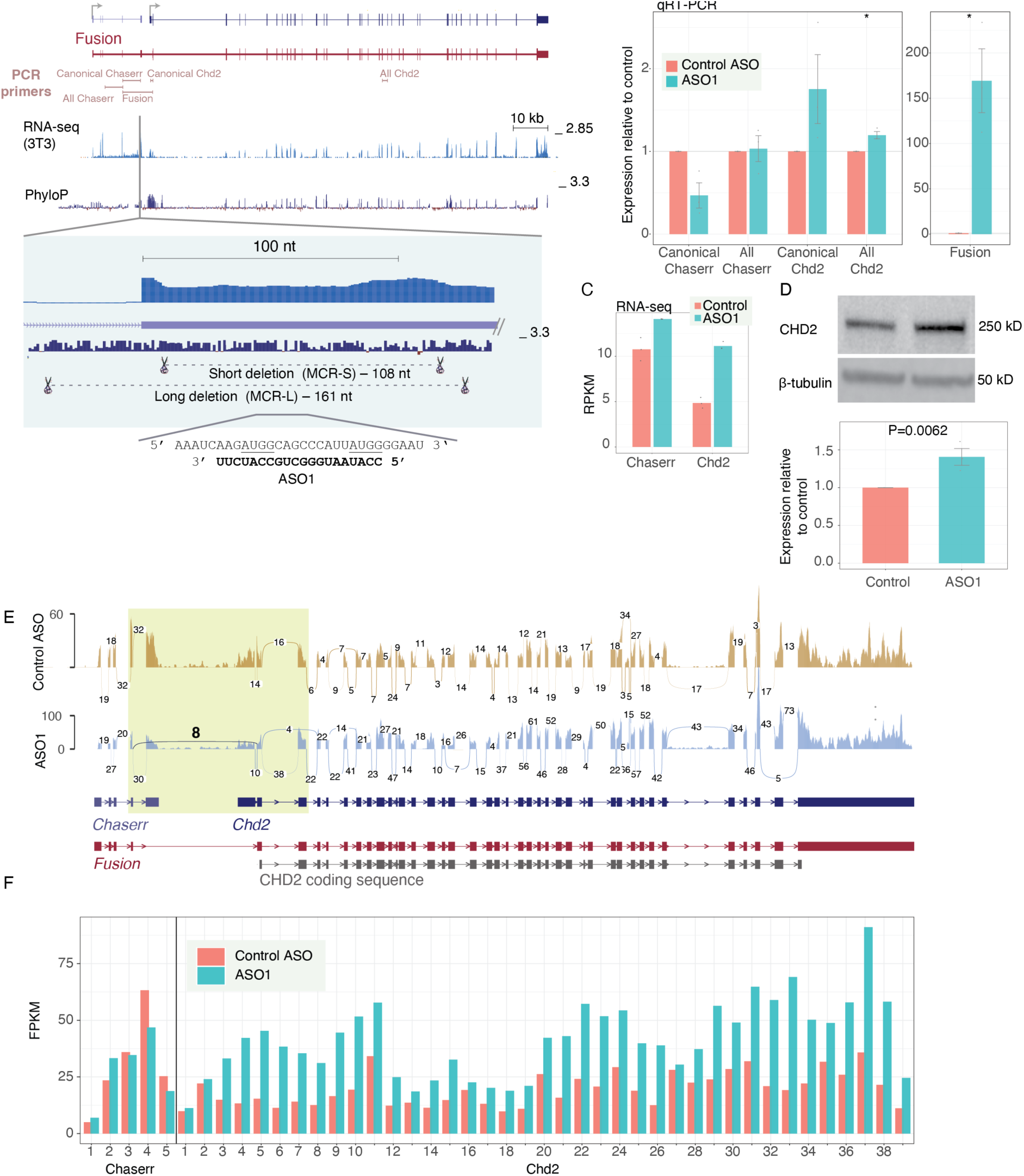
Treatment with ASO1 results in the formation of a Chaserr-Chd2 fusion transcript. **(A)** Overview of the *Chaserr*-*Chd2* locus in mouse, showing RNA-seq read coverage in unperturbed 3T3 cells. Positions of pairs of primers used for qRT-PCR are shown. PhyloP conservation based on a 100- mammal whole genome alignment is taken from the UCSC genome browser. ASO1 binding site and the regions deleted in the mouse models are shown at the bottom. **(B)** qRT-PCR quantification of the indicated transcripts following ASO1 or control ASO transfection into 3T3 cells for 48h. Expression is normalized to β-actin, *n* = 3 * - P<0.05, two-sided t-test. **(C)** RNA-seq data quantification for Chaserr and Chd2, in 3T3 cells treated with the indicated ASOs, n = 3. **(D)** Western blot of CHD2 expression in 3T3 cells treated with either ASO1 or control ASO for 48h, n = 3 . **(E)** Sashimi plot indicating the numbers of reads spanning individual splice junctions in *Chaserr* and *Chd2* in 3T3 cells treated with the indicated ASOs. Only junctions supported by more than 3 reads in the aggregated data are shown. The annotated main isoforms of Chaserr and Chd2, the estimated fusion isoform, and the canonical CHD2 coding sequence are shown below. The fusion splice junction is highlighted **(F)** RNA-seq based estimation of expression levels of individual exons in *Chaserr* and *Chd2*, in 3T3 cells treated with the indicated ASO. The plot indicates the average expression per exon calculated over three replicates, as the number of fragments per exon per kilobase per million mapped reads (FPKM).

## Results

### Treatment with Chaserr MCR-targeting ASOs leads to the formation of a Chaserr-Chd2 fusion transcript

To study the functional consequences of targeting the RRAUGG motifs in the *Chaserr* MCR, we profiled NIH3T3 (‘3T3’) cells transfected with ASO1 (**Fig. 1A**) using qRT-PCR, RNA-seq, and Western blot (WB). ASO1 led to a ∼2-fold increase in *Chd2* mRNA levels and up-regulated *Chaserr* levels (**Fig. 1B-C**), consistent with our previous observations in Neuro2a cells^10^. As expected, this increase in *Chd2* mRNA resulted in an increase in CHD2 protein (**Fig. 1D**). We then inspected the coverage of RNA-seq reads at the *Chaserr*-*Chd2* locus. In addition to the regular spliced reads covering *Chaserr* and *Chd2* exons, we detected an unannotated splice junction between the penultimate exon of *Chaserr* and the second exon of *Chd2* that was strongly induced in ASO1-treated cells (**Fig. 1E**). This was accompanied by a decrease in the number of reads spanning the canonical last splice junction of *Chaserr*, indicating that a portion of *Chaserr* transcription events produced a *Chaserr*–*Chd2* fusion transcript instead of producing canonical *Chaserr* transcripts (**Fig. 1E**). We further quantified *Chd2* exon coverage and observed an average increase of 2.7-fold across exons 2–39, with no change in the coverage of the first exon — suggesting that the formation of the *Chaserr*–*Chd2* fusion did not come at the expense of the canonical isoform (**Fig. 1F**). Aside from fusion formation, no other notable changes in splicing patterns were observed. Thus, the fusion transcript appears to contain the full sequence from *Chaserr* exons 1–4 and *Chd2* exons 2–39, forming a chimeric mRNA molecule that encodes the full 5,481 nt ORF of *Chd2* preceded by an alternative 5’ UTR (‘fusion 5’ UTR’). Specifically, this fusion 5’ UTR is composed of exons 1–4 of *Chaserr* and the beginning of Chd2 exon 2 (**Fig. 1A**).

To validate our observations from RNA-seq, we designed qRT-PCR primers to specifically detect the fusion splice junction. We found that, although detectable at baseline levels in cells treated with the control ASO, treatment with ASO1 significantly increased fusion levels while simultaneously reducing canonical Chaserr by ∼2-fold (**Fig. 1A-B**). We note that the very low, yet detectable, levels of the fusion junction in untreated cells, or in those transfected with the control ASO, make it difficult to determine the exact fold-change of its increase. Primers spanning the canonical first intron of *Chd2* did not identify a reduction in the expression of the canonical Chd2 exon1-exon2 splice junction, further supporting the notion that fusion formation does not impact *Chd2* transcription (**Fig. 1B**). To validate that the formation of the fusion transcript depends on continuous transcription in the intergenic region between *Chaserr* and *Chd2*, we transfected 3T3 cells with ASO1 or a control ASO alongside GapmeRs targeting *Chaserr*, *Chd2*, or the intergenic region (**Fig. S1A-B**). As predicted, the GapmeR that more effectively reduced *Chaserr* expression (targeted to a region close to its end), the two GapmeRs that target the intergenic region, and the *Chd2*-targeted GapmeR all strongly reduced the formation of the fusion transcript in ASO1-treated cells (**Fig. S1B**).

To test for the conservation of this phenomenon, we designed hASO1, which targets the region in the human CHASERR MCR that is aligned to the region targeted by ASO1 in mice (**Fig. S2A**). This 20-nt ASO targets two consecutive UAGG motifs conserved between human and mouse, with three nucleotide differences in the short stretch between the two motifs (**Fig. S2B**). Treatment of the MCF7 breast cancer cell line led to a strong induction of a corresponding *CHASERR*–*CHD2* fusion transcript as quantified by qRT-PCR (**Fig. S2C**). This was supported by an increase in fusion-specific RNA-seq reads, along with an increase in total CHD2 expression (**Fig. S2D-E**) and a corresponding increase in CHD2 protein (**Fig. S2F**). ASOs targeting the *CHASERR* MCR thus induce *Chaserr*-*Chd2* fusion transcript formation in both mouse and human cells.

### Deletion of Chaserr MCR regions induces the formation of the *Chaserr*-*Chd2* fusion transcript

To examine the consequences of *Chaserr* MCR disruption using an orthogonal approach and in an organismal setting, we used CRISPR-Cas9 with several gRNAs targeting the Chaserr MCR region and obtained transgenic mice with either a short (*Chaserr*^MCR-S^ allele) or a longer deletion at the beginning of the last exon of Chaserr (*Chaserr*^MCR-L^ allele, **Fig. 1A**). The *Chaserr*^MCR-S^ allele lacks 108 nt that correspond to bases 10–117 in the last *Chaserr* exon whereas *Chaserr*^MCR-L^ allele has a deletion of 161 nt including the last 34 nt of the last intron and the first 127 nt of the last exon. The deletions were confirmed by PCR (**Fig. S2A**). Both heterozygous and homozygous mice carrying the *Chaserr*^MCR-S^ and *Chaserr*^MCR-L^ alleles were born close to the expected Mendelian ratios (**Fig. S2B**), with only a minor reduction in the number of *Chaserr*^MCR-S/MCR-S^ pups. This suggests that the *Chaserr* MCR loss is less consequential for mouse viability than the complete loss of *Chaserr* transcription in the promoter-deletion *Chaserr^−/−^* mice^7^. Loss of MCR resulted in a strong increase in fusion transcript formation as quantified by qRT-PCR while having minimal effects on reads spanning the first *Chd2* intron (**Fig. 2B-D**, **S2C**). RNA-seq analysis showed that *Chaserr*^MCR-L/MCR-L^ mice also had a substantial retention of the last intron of *Chaserr*, likely because it lacked the 3’ splice site (**Fig. 2A**), and so we focused further analysis on the MCR-S mice, which exhibited unaltered splicing. In adult whole-brain tissue, *Chd2* mRNA and protein were increased in *Chaserr*^MCR-S/MCR-S^ mice with dose-sensitive effects (**Fig. 2E-F**, **S2D**). RNA-seq read coverage was uniformly increased in exons 2–39 of *Chd2* (**Fig. 2F**). Loss of the Chaserr MCR thus leads to *Chaserr*-*Chd2* fusion transcript formation, and increased CHD2 protein levels also *in vivo*, without the strong effect on mouse viability caused by the *Chaserr* promoter deletion.

**Figure 2.**
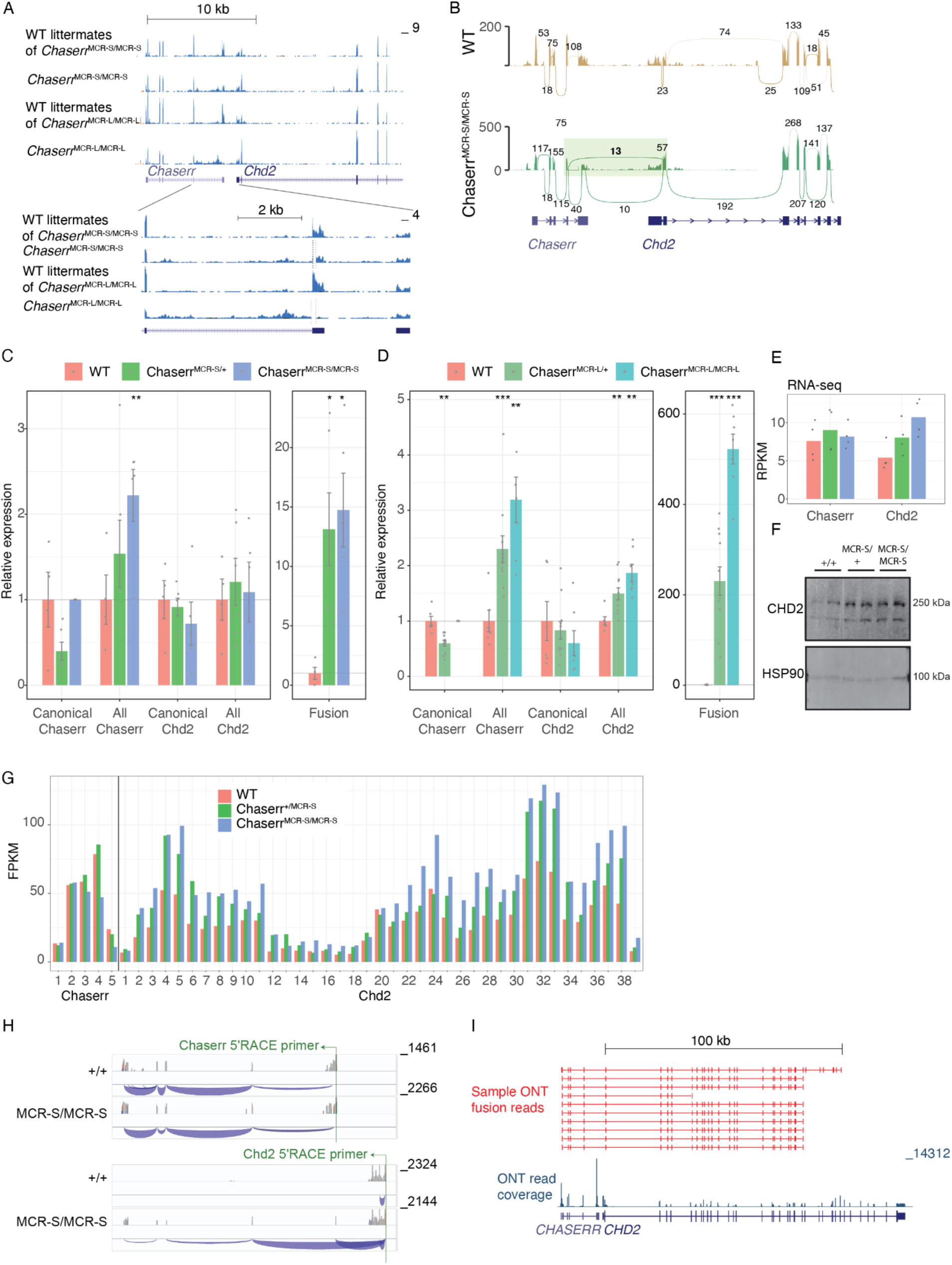
Deletion of the Chaserr MCR leads to Chaserr-Chd2 fusion transcript formation. **(A-B)** RNA-seq based read coverage (A) and counts of splice-site–spanning reads (B) in the whole brain of adult mice with the indicated genotypes. Numbers on the arcs indicate raw read counts. Fusion junction region is shaded. **(C-D)** Quantification of the indicated transcripts by qRT-PCR on RNA from the whole brain of adult mice with the indicated genotypes. P-values computed using two-sided t-test. * - P<0.05, ** - P<0.005, *** - P<0.0005. Canonical *Chaserr* can not be quantified in homozygous animals as they lack the region targeted by the PCR primers. **(E)** RNA-seq–based transcript quantification. **(F)** Western blots of protein from whole brains of mice with the indicated genotypes and genders. **(G)** Per-exon RNA-seq read coverage in aggregated data from the indicated genotypes, calculated as number of reads per exon per kilobase per million mapped reads, averaged over four biological replicates. **(H)** 5’ RACE from the indicated primer followed by RNA-seq in the whole brain from the indicated genetic background. Read coverage and splice junctions are shown. **(I)** Oxford Nanopore long-reads from ^12^ mapped to the human genome. Representative long reads supporting the fusion transcript are shown.

### The *Chaserr*-*Chd2* fusion mRNA is efficiently exported from the nucleus and is an NMD substrate

The uniform increase in expression of *Chd2* exons in the ASO1 treatment or MCR deletion models with no notable changes in splicing patterns except for fusion formation suggest the fusion transcript contains the full sequence from *Chaserr* exons 1–4 and *Chd2* exons 2–39. Supporting full transcript connectivity, 5’ RACE on RNA from mouse brain using a primer located in *Chd2* exon 2 recovered a signal from the first exon of *Chaserr* (**Fig. 2H**). We also found long reads corresponding to full-length fusion transcripts in nanopore sequencing reads from human cells^12^ (**Fig. 2I**).

To profile the fate of the fusion mRNA, we first performed subcellular fractionations in 3T3 cells treated with control or ASO1 to induce fusion formation, followed by qRT-PCR to quantify the localization of the canonical *Chaserr* and *Chd2*, and their fusion (**Fig. 3A**). Fusion RNA was found in the cytoplasmic fraction at levels similar to the canonical *Chd2* mRNA, which was overall more nuclear than *Chaserr*, possibly because of its length, and in agreement with our previous observations in mESCs^7^. Consistent with the nuclear export of the fusion transcript, we observed occasional co-localization of signals arising from probes targeting *Chaserr* and *Chd2* transcripts in the cytoplasm of ASO1-treated 3T3 cells (**Fig. 3B**). We previously found that *Chaserr* is strongly targeted by nonsense-mediated decay (NMD)^7^, which degrades transcripts with translated ORFs ending upstream of the last splice junction. To determine whether the fusion transcript undergoes NMD we examined the sensitivity of *Chaserr*, *Chd2*, and the fusion to cyclohexamide (CHX), which inhibits translation and alleviates NMD. CHX treatment in 3T3 cells led to increased expression of all three transcripts, and increased fusion transcript levels in cells treated with control ASO or ASO1 (**Fig. 3C**), suggesting that *Chaserr*, *Chd2*, and their fusion are all NMD substrates to varying degrees. Similar results were obtained for the human transcripts (**Fig. S3A-B**).

**Figure 3.**
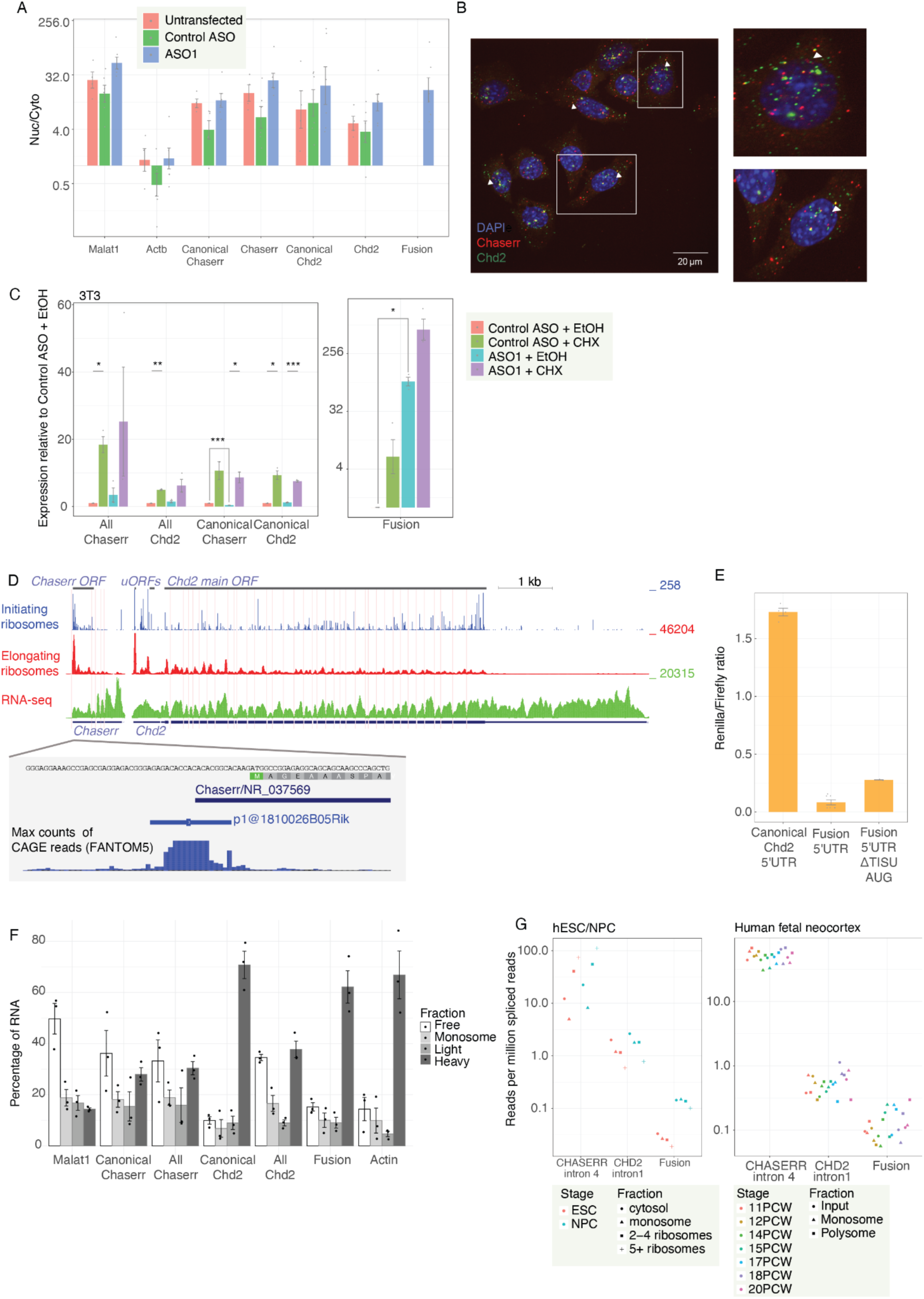
Translation of the Chaserr-Chd2 fusion mRNA. **(A)** qRT-PCR of 3T3 cells, measuring the nuclear/cytoplasmic (Nuc/Cyto) ratios of the indicated genes following the indicated treatments. **(B)** HCR image showing localization of Chaserr and Chd2 in ASO1-treated 3T3 cells. Arrows indicate colocalized Chaserr and Chd2 probe signals. **(C)** qRT-PCR in 3T3 cells measuring the indicated transcripts treated for 48 hr with the indicated ASO followed by an 8h treatment with Cyclohexamide (CHX) or its EtOH vehicle. Relative expression normalized to β-actin, n = 3. **(D)** Alignment of Ribo-seq and RNA-seq reads from aggregated mouse experiments in GWIPS-VIZ ^13^. The longest ORF in Chaserr, and the two longest uORFs in the Chd2 5’UTR are indicated. Ribo-seq reads are separated into those from regular Ribo-seq libraries sequencing footprints of elongating ribosomes, and ribosomes treated with reagents that arrest them at translation initiation sites. **(E)** Luciferase assay comparing luminescence from the indicated 5’UTR fused to Renilla luciferase to a control Firefly luciferase expressed from the same plasmid. (**F**) Polysome profiling of ASO1-treated 3T3 cells. RNA distribution across polysome fractions (free, monosome, light and heavy) is shown for each gene. Cells were profiled 48 hr after transfection with ASO1. **(G)** Normalized RNA-seq read counts for the corresponding splice junctions in the indicated fraction of cells from the indicated sample. Data from ^14^ (left) and from ^15^ (right).

### The *Chaserr*-*Chd2* fusion mRNA is translated

The fusion 5’ UTR is 629 nt, comparable in length to the 571 nt of the canonical *Chd2* 5’UTR (**Fig. 3D**). Similarly, the human fusion 5’ UTR is 640 nt in comparison to the 572 of the canonical *CHD2* 5’ UTR. *Chaserr* exons 1–4 are more G/C rich than the canonical Chd2 5’UTR (63% G/C vs. 44% G/C, respectively), which contains an upstream ORF (uORF) of 354 nt in mouse and 126 nt in human. In comparison, the canonical *Chd2* 5’ UTR contains only short uORFs <40 nt in length. The CHASERR ORF starts with a CAATAUGGCCGG sequence, which matches the first ten bases of the SAASAUGGCGGC (S=C/G) TISU motif found in some mRNAs with particularly short 5’UTRs^17,18^ and is also highly conserved at the 5’ of the *Cyrano* lncRNA^10^. The ORF in *Chaserr,* and the uORFs in the *Chd2* 5’ UTR all show evidence for translation in human and mouse Ribo-seq datasets (**Fig. 3D** and **Fig. S3C**).

To test how these features control the efficiency of translation, we cloned the sequences of three possible mouse *Chd2* 5’ UTRs: the canonical *Chd2* 5’ UTR, the fusion 5’ UTR and a fusion 5’ UTR where the TISU element was deleted upstream of the Renilla luciferase in a psiCheck2 backbone expressing the Firefly luciferase from a separate promoter. When transfected into 3T3 cells, the Renilla mRNA with the fusion 5’ UTR was translated, but at levels ∼20-fold lower than an mRNA containing the canonical 5’ UTR. Deletion of the TISU element, including the start codon of the uORF, led to a 2-fold increase in luciferase levels compared to the wild-type fusion 5’ UTR, suggesting that this conserved element may also control the translation of CHD2 from the fusion transcript (**Fig. 3E**). We note that subtle variation in the transcription start site of *Chaserr* can affect the inclusion of the uORF and thus differentially affect translation from the fusion mRNA (see Discussion). We further tested the role of this ORF by treating 3T3 and MCF7 cells with an ASO targeted to the TISU element. This led to an overall increase in *Chaserr* and fusion RNA in both cell lines, suggesting that the uORF may regulate the susceptibility of both transcripts to NMD (**Fig S3D-E**).

To profile the translation of the fusion mRNA expressed from the genome, we used qRT-PCR on polysome fractions of ASO1-treated 3T3 cells. Fusion transcripts were found in the polysomal fraction at levels comparable to those of the endogenous *Chd2* (**Fig. 3F** and **S3F**). We also analyzed data from two studies that used RNA sequencing on polysomal fractions from human embryonic stem cells as well as from neuronal progenitors derived from them^14^, and from the human fetal neocortex^15^. Comparison of normalized read counts spanning *Chaserr*-, *Chd2-*, and fusion-specific splice junctions showed that the fusion is found on polysomes with frequencies comparable to those of the canonical *Chd2* and that its levels are increased in the neuronal lineage and during fetal neurogenesis (Pearson’s R=0.33, P=0.03 between post-conception week and normalized fusion-specific reads across all samples, **Fig. 3H**) reaching levels on heavy polysomes comparable to those of canonical *Chd2* in the later-stage embryonic development (**Fig. 3H**). We conclude that the fusion transcript is translated at lower levels than the canonical Chd2 mRNA, most likely because *Chaserr* uORF sequence and/or its G/C rich first exon inhibit ribosomal scanning from reaching the *Chd2* start codon; however, translation of the fusion can be efficient, particularly in a neuronal context.

### Perturbations of *Chaserr’s* last splice site and polyadenylation sites up-regulate Chaserr- Chd2 fusion

Since the Chaserr MCR lies close to the beginning of the last exon, we hypothesized that it controls the splicing of this last exon, and that this is coupled to *Chaserr* cleavage and polyadenylation (CPA). We, therefore, sought *cis*-acting sequences and *trans*-acting factors that cooperate with the MCR in regulating fusion transcript formation.

To identify *trans*-acting protein-coding genes that regulate *Chaserr*-*Chd2* fusion formation, we mined the ENCODE resource of RNA-seq datasets from K562 and HepG2 cells where shRNAs were used to deplete different RNA-associated factors^19^. In each of the two cell lines, we mined perturbations that increase the formation of the *Chaserr*-*Chd2* fusion by querying for RNA-seq reads containing the specific fusion junction and comparing them to the number of reads covering the last *Chaserr* exon-exon junction and the first *Chd2* exon-exon junction (as the latter two are not formed when the fusion transcript is formed). While most perturbations had no effect, shRNAs targeting specific proteins led to an increase in the number of reads emanating from the *Chaserr*-*Chd2* fusion (**Fig. 4A-B**). These included *CPSF6*, which regulates a subset of polyadenylation events^20^, *PUF60* which is a canonical splicing factor^21^, and *NSUN2*, a methyltransferase that catalyzes the 5-methylcytosine (m5C) RNA modification. We analyzed the available data on m5C modifications but found no evidence for its specific deposition in *Chaserr* or *Chd2*. Knockdown (KD) of *CPSF6* and *PUF60* increased fusion formation in human MCF7 cells, whereas *NSUN2* KD did not have an effect (**Fig. 4C**). Notably, PUF60 KD also led to increased levels of *CHASERR* (**Fig 4C**). We analyzed the available CLIP data (available for CPSF6 in human^19^ and PUF60 in mouse^22^) and identified binding sites for *CPSF6* and *PUF60* near the poly(A) signal of *Chaserr* and near the 2^nd^ exon of Chd2 (**Fig. 4D**).

**Figure 4.**
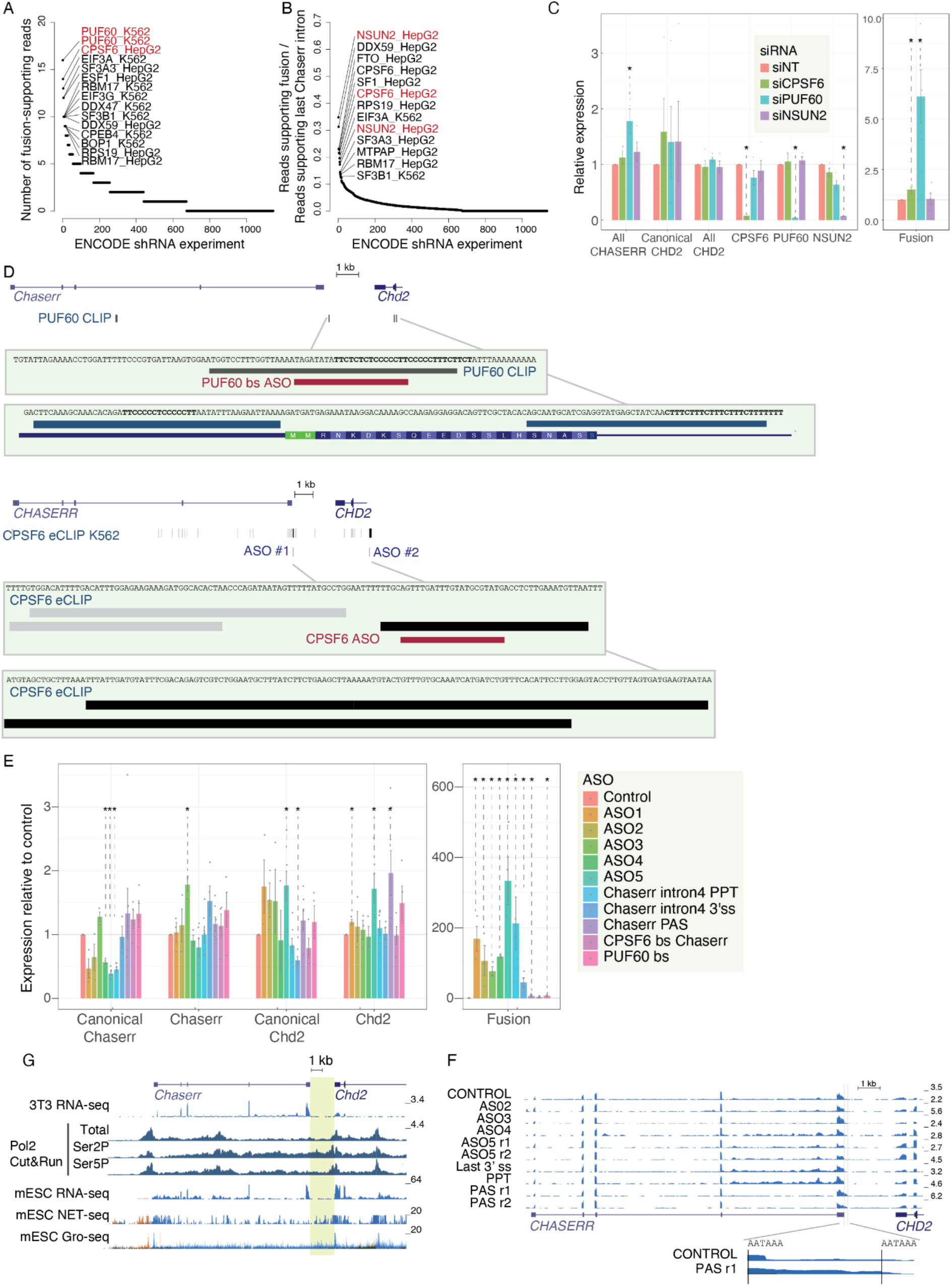
Trans factors and cis-acting sequences modulate fusion transcript formation. **(A)** ENCODE shRNA-RNA-seq experiments that show evidence of Chaserr-Chd2 transcript formation. Experiments are ranked by the number of reads containing a 20mer sequence supporting the specific fusion junction. **(B)** As in A, showing the ratio between the number of reads supporting the specific Chaserr-Chd2 and the number of reads supporting the splicing of intron 4 of canonical Chaserr. **(C)** qRT-PCR for the indicated genes in MCF7 cells. * - P<0.05, two-sided t-test. **(D)** Top: CLIP data from mouse cells for PUF60 and the corresponding ASO position. Bottom: CLIP data for CPSF6 in human cells (color-coded based on the ENCODE pipeline with darker colors corresponding to more confident sites) and the corresponding ASO position. **(E)** qRT-PCR for the indicated transcripts in 3T3 cells treated with the indicated ASOs.* - P<0.05, two-sided t-test. **(F)** RNA-seq read coverage in 3T3 cells treated with the indicated ASO (r1/r2 - biological replicates). The region around the Chaserr 3’ end is magnified and positions of the AAUAAA PAS motifs are indicated. **(G)** Evidence for continuous transcription through the Chaserr-Chd2 intergenic region. Shown is data for RNA-seq and Cut&Run from 3T3 cells using antibodies for total, Ser2-phosphorylated and Ser5-phosporylated Pol2 (this study); RNA-seq and NET-seq from mouse embryonic stem cells (mESCs) from^23^; GRO-seq data from mESCs from ^24^. NET-seq and GRO-seq read count was capped at 20. The intergenic region between Chaserr and Chd2 is shaded. For the RNA data, blue indicates coverage on the – strand of the genome (that *Chaserr* and *Chd2* are transcribed from) and red indicates coverage on the + strand.

To identify *cis*-acting sequences in the *Chaserr* locus that may regulate fusion transcript formation, we designed a series of ASOs targeting regions potentially involved in splicing and polyadenylation of the last exon of *Chaserr*. These ASOs blocked the polypyrimidine tract (PPT) or the 3’ splice site of the last intron of *Chaserr*, conserved sequences preceding the MCR (ASO4 and ASO5), the polyadenylation signal (PAS), and binding sites of two RNA binding proteins CPSF6 and PUF60 in the vicinity of the *Chaserr* poly(A) site. We also evaluated ASO2 and ASO3, two previously designed ASOs^10^ that target additional RRUAGG motifs in the last *CHASERR* exon (**Fig. S1A**). Importantly, the target sites of ASO2 and ASO3 are located 72 and 167 nt from the start of the exon — providing a valuable readout for targeting conserved motifs that are positioned substantially away from both the 3’ splice site and the PAS (**Fig. 4D**). When transfected into 3T3 cells, all the ASOs caused substantial increases in the formation of the *Chaserr*-*Chd2* fusion as evaluated by qRT-PCR and RNA-seq (**Fig. 4E**). The ASOs had variable effects on canonical *Chaserr* and *Chd2* transcripts. As expected, the ASOs targeting the 3’ splice site adjacent regions reduced the efficiency of the splicing of the last *Chaserr* intron.

Those targeting the polyadenylation signal shifted the canonical poly(A) site by ∼140 nt to a downstream AAUAAA element (**Fig. 4F**). Similar results were obtained in human MCF7 cells (**Fig. S4A**). These results suggest that efficient splicing is required for the *CHASERR* CPA and the termination of its transcription. As such, blocking of these sites likely results in continued Pol2 elongation across the full length of the *Chd2* locus, providing an opportunity for the formation of the splice junction between the 4^th^ exon of *Chaserr* and the 2^nd^ exon of *Chd2*. This response is also triggered by targeting RRUAGG motifs that are several hundreds of bases away from both the splice site and the PAS - thus transcription termination is also dependent on specific sequence elements in the last exon of *Chaserr*. Inspection of nascent RNA-seq indicates that Pol2 continues to transcribe across the intergenic region under native conditions (**Fig. 4G**). Therefore, it is plausible that the ASOs merely prevent cleavage of *Chaserr* and do not increase Pol2 occupancy that may interfere with transcription initiation of canonical *Chd2* mRNA.

We conclude that various sequences and trans-acting RNA binding proteins orchestrate the proper splicing and CPA of the last exon of *Chaserr*, and blocking perturbations at those sequences or depleting the *trans*-acting factors interacting with them leads to increased formation of the *Chaserr*-*Chd2* fusion.

### *Chaserr*-*Chd2* fusion is induced by neuronal activation and neuronal gene expression is affected by MCR loss

Following the observation that alternative usage of *Chaserr* as a 5’ UTR can affect translation of the *CHD2* mRNA, we hypothesized that expression of the *Chaserr*-*Chd2* fusion may be induced under specific biological conditions, in particular in the neuronal context, where fusion appears well translated (**Fig. 3G**). To explore this, we mined various neuronal RNA-seq datasets available on SRA. We found that the fusion transcript is substantially induced during neuronal activation in mouse cortical neurons and fetal human neurons^25^ (**Fig. 5A-B**). Interestingly, we found that the predominant fusion isoform induced during activation of mouse neurons consists of a splice junction that spans from the middle of the fifth *Chaserr* exon, and thus the predicted 5’UTR contains five exons from Chaserr and the 2^nd^ exon of *Chd2*. This isoform is notably not induced with ASO1 treatment, and this splice site is not conserved in human Chaserr (which induces the ‘regular’ fusion transcript upon neuronal activity). We term this *Chaserr*-*Chd2* fusion isoform the ‘mouse-specific fusion’ (**Fig. 5C**).

**Figure 5.**
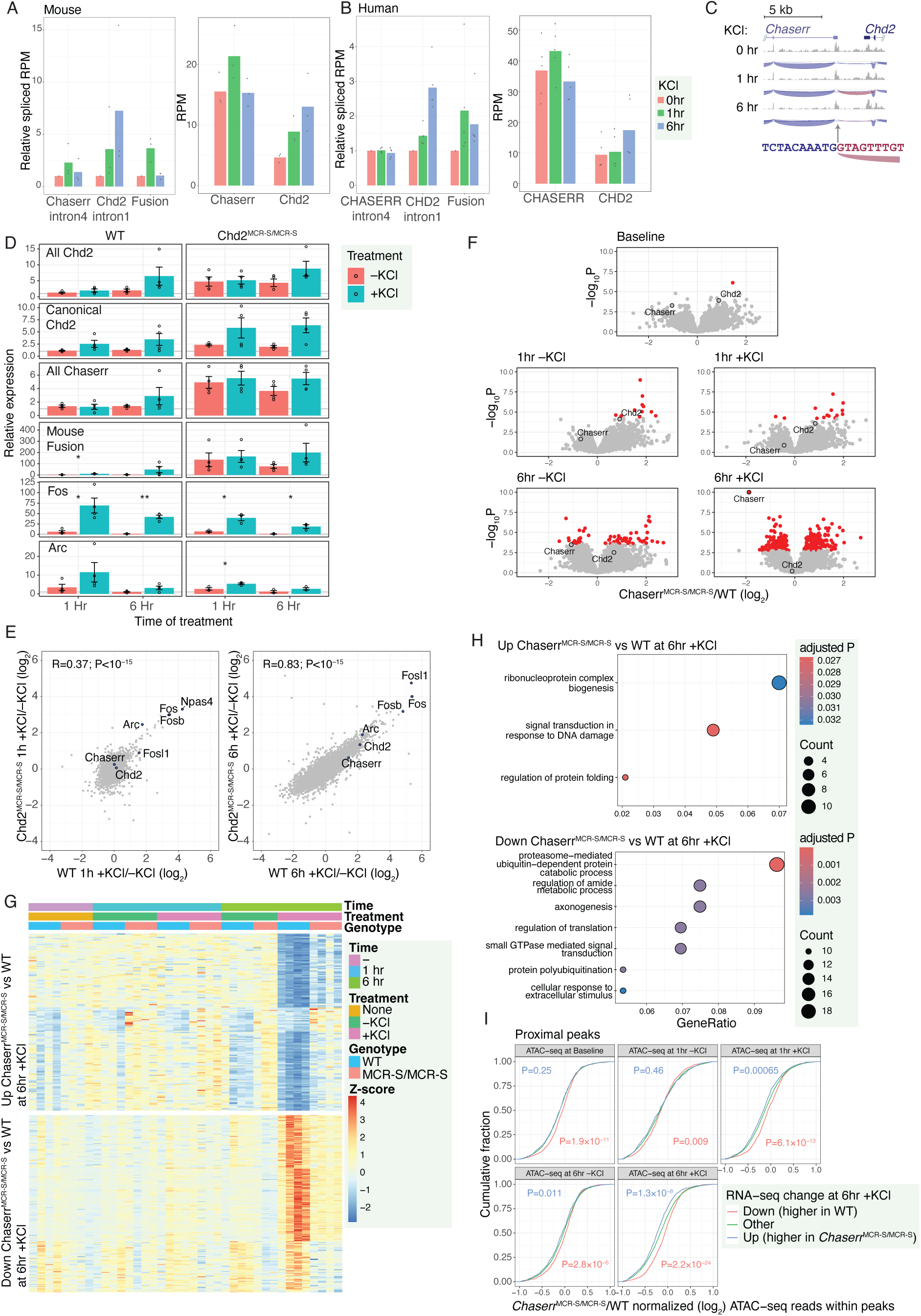
*Chaserr*^MCR-S/MCR-S^ neurons exhibit dysregulated neuronal activity-induced gene expression and chromatin accessibility. **(A-B)** Left: Splice-junction–spanning reads for the indicated introns (‘Chaserr intron4’ is found only in canonical *Chaserr*, and ‘Chd2 intron1’ only in canonical *Chd2*), normalized to the 0hr time point; Right: FPKM RNA-seq quantification; at the indicated time after KCl treatment, in mouse primary cortical neurons (A), and human fetal brain cultures (B), both from ^25^. (C) RNA-seq read coverage and intron-spanning reads in the mouse RNA-seq data, indicating the position of the mouse-specific 5’ splice site in the last exon of *Chaserr*, with the sequence of the canonical 5’ splice site shown. **(D)** qRT-PCR measurements of the indicated transcripts at the indicated time point after KCl treatment of primary mouse cortical neurons. * - P<0.05; ** - P<0.005, two-sided t- test comparing –KCl and +KCl samples. **(E)** Correspondence between changes in gene expression of WT and *Chaserr*^MCR-S/MCR-S^ primary cortical neurons following 1 hr (left) or 6 hr (right) of KCl treatment. Spearman’s correlation and P-values are shown, and *Chaserr*, *Chd2*, and select immediate-early genes are highlighted. **(F)** Volcano plots for changes in gene expression when comparing WT and *Chaserr*^MCR-S/MCR-S^ neurons with the indicated treatment. Fold-changes and P values computed using DESeq2 ^27^. Genes with adjusted P<0.05 are in red. **(G)** Scaled gene expression patterns of the genes up-regulated (top) and down-regulated (bottom) in *Chaserr*^MCR-S/MCR-S^ primary cortical neurons compared to WT controls 6hr +KCl samples. Each row corresponds to a gene scaled to mean 0 and standard deviation of 1. **(H)** GO enrichment computed using the clusterProfiler package after simplification for the genes shown in G. **(I)** Changes in accessibility in ATAC-seq data from the indicated samples, comparing the peaks found within 2kb of the transcription start sites of the genes shown in G and other genes. P-values computed using the Wilcoxon two-sided rank sum test comparing the peaks associated with the up- and down-regulated genes compared to other genes are shown.

To investigate the connection between neuron depolarization and fusion transcript formation, we cultured primary cortical neurons from WT and *Chaserr*^MCR-S/MCR-S^ mice for 7 days in vitro. The cells were then mock-treated or depolarized with 55 mM KCl for 1 or 6 hours. After treatment, we profiled gene expression changes by qPCR and RNA-seq. The fusion transcript increased substantially in KCl-treated WT neurons, reaching a 50-fold increase at six hours of activation, with expression levels comparable to uninduced *Chaserr*^MCR-S/MCR-S^ neurons (**Fig. 5D**). In comparison, only a marginal increase in fusion was observed in activated *Chaserr*^MCR-S/MCR-S^ neurons, where fusion was already well-expressed at baseline. In both the WT and *Chaserr*^MCR-S/MCR-S^ activated neurons, the increase in fusion was accompanied by an increase in both *Chaserr* and canonical *Chd2*, suggesting that fusion induction is associated with increased transcription across the locus.

To study the effects of constitutive production of the fusion transcript on neuronal activity, we profiled the neurons by RNA-seq. In both genotypes, KCl treatment resulted in substantial changes in gene expression and induction of the canonical immediate early genes such as *Fos*, *Arc*, and *Npas4* (**Fig. 5D,E**). While few genes changed in expression at baseline conditions, pronounced changes in gene expression appeared in cells treated with KCl for 6 hr, with 153 genes being up- and 202 genes down-regulated by at least 25%, respectively, with adjusted P<0.05 (**Fig. 5F**). Roughly half of the genes affected in the 6hr +KCl sample were also affected in a similar direction at 1hr or at baseline, albeit to a lesser degree and the other half was affected specifically in the 6 hr activated neurons, where these genes were substantially less activated or less repressed in *Chaserr*^MCR-S/MCR-S^ cells (**Fig. 5G**). Gene Ontology analysis found several categories potentially related to neuronal activity in the differentially expressed genes, such as DNA damage, known to be an important part of transcriptional regulation during neuronal activation^26^, enriched in the up-regulated genes and axonogenesis and translational control enriched in the down-regulated ones (**Fig. 5H**).

Since CHD2 is a chromatin remodeler, we wondered if the changes in gene expression are associated with changes in chromatin accessibility. To test this, we used ATAC-seq under the same conditions in cultured WT and *Chd2*^MCR-S/MCR-S^ neurons. Collectively, the promoters of differentially expressed genes exhibited differential accessibility in *Chd2*^MCR-S/MCR-S^ neurons (**Fig. 5I**), with the strongest effects observed in promoters of genes that were down-regulated in *Chd2*^MCR-S/MCR-S^ neurons. Surprisingly, these down-regulated genes were significantly more accessible in *Chd2*^MCR-S/MCR-S^ neurons under baseline conditions, and even more so following KCl-induced activation. We conclude that the constitutive expression of *Chaserr*-*Chd2* fusion transcripts in *Chd2*^MCR-S/MCR-S^ neurons is accompanied by differential chromatin accessibility, which has the most profound consequences on gene expression when neurons are activated.

### ASO1 fusion induction rescues CHD2 expression and behavioral phenotypes in a mouse model of CHD2 haploinsufficiency

The targeted blocking of conserved motifs in the last exon of *Chaserr* thus produces an additional template for full-length CHD2 protein without substantially affecting the expression of CHD2 from the canonical *Chd2* mRNA. This introduces a new strategy for treating haploinsufficient disorders, through modulation of the transcription termination process of an upstream gene transcribed in close proximity and on the same strand. Such disorders are common as >1500 human genes are estimated to be haploinsufficient ^28^ and >170 of those are associated with known disorders in ClinGen ^29^. To demonstrate the therapeutic potential of the fusion-inducing ASOs we used the *Chd2*^+/m^ mouse model that was established by our lab^7^. On the 129X1/SvJ background, these mice recapitulate key phenotypes observed in CHD2 patients, including motor and cognitive deficits^30^.

To study the ability of ASO1 to affect CHD2-mediated neurological phenotypes, we used *Chd2*^+/m^ mice bred to the 129X1/SvJ for at least 10 generations. Since the phenotypes described below were most pronounced in relatively young mice, we administered perinatal intracerebroventricular (ICV) injections of ASO1 or a control ASO to WT and *Chd2*^m/+^ mice at postnatal day 2 (P2). We first validated the broad distribution in the brain by using FAM-labelled ASOs (**Fig. S5A**). We then evaluated the effect of both the *Chd2*^+/m^ genotype and ASO injection on three robust phenotypes exhibited by the *Chd2*^+/m^ mice: the hindlimb clasping, hanging wire grip, and the Y-maze tests at the age of 4–5 weeks. Under normal conditions, when held upside down, mice will extend their hindlimbs outward from the abdomen, displaying typical motor control and reflexes. In contrast, mice with neurological impairments often exhibit a clasping behavior, pulling their hindlimbs tightly towards their abdomen. Clasping scores were higher in *Chd2*^+/m^ mice compared to WT littermates, and the ASO had a significant corrective effect on this phenotype associated with motor coordination and neurological function (**Fig. 6A-B**). In the hanging wire grip test, mice were placed on a wire by their forelimbs and allowed to grasp the wire with their hindlimbs to stabilize themselves. *Chd2*^+/m^ mice demonstrated a significant reduction in hanging time duration, exhibited decreased stability on the wire, and fell more frequently than WT mice, with the phenotypes reversed by ASO injection (**Fig. 6C-E**). In the third test, we used a Y-maze consisting of three arms of equal length and angle. The mice were initially allowed to explore just two of the three arms and then reintroduced to the maze for an additional 5 minutes. The *Chd2*^+/m^ mice entered the novel arm significantly fewer times and spent less time there, indicating potential cognitive impairments that were reversed by ASO1 injection (**Fig. 6F-H**). These behavioral assays collectively suggest that heterozygous loss of CHD2 in mice leads to observable deficits in motor coordination, neuromuscular strength, and cognitive function and that these can be at least partially rescued by the *Chaserr*-targeting and fusion-inducing ASO1.

**Figure 6.**
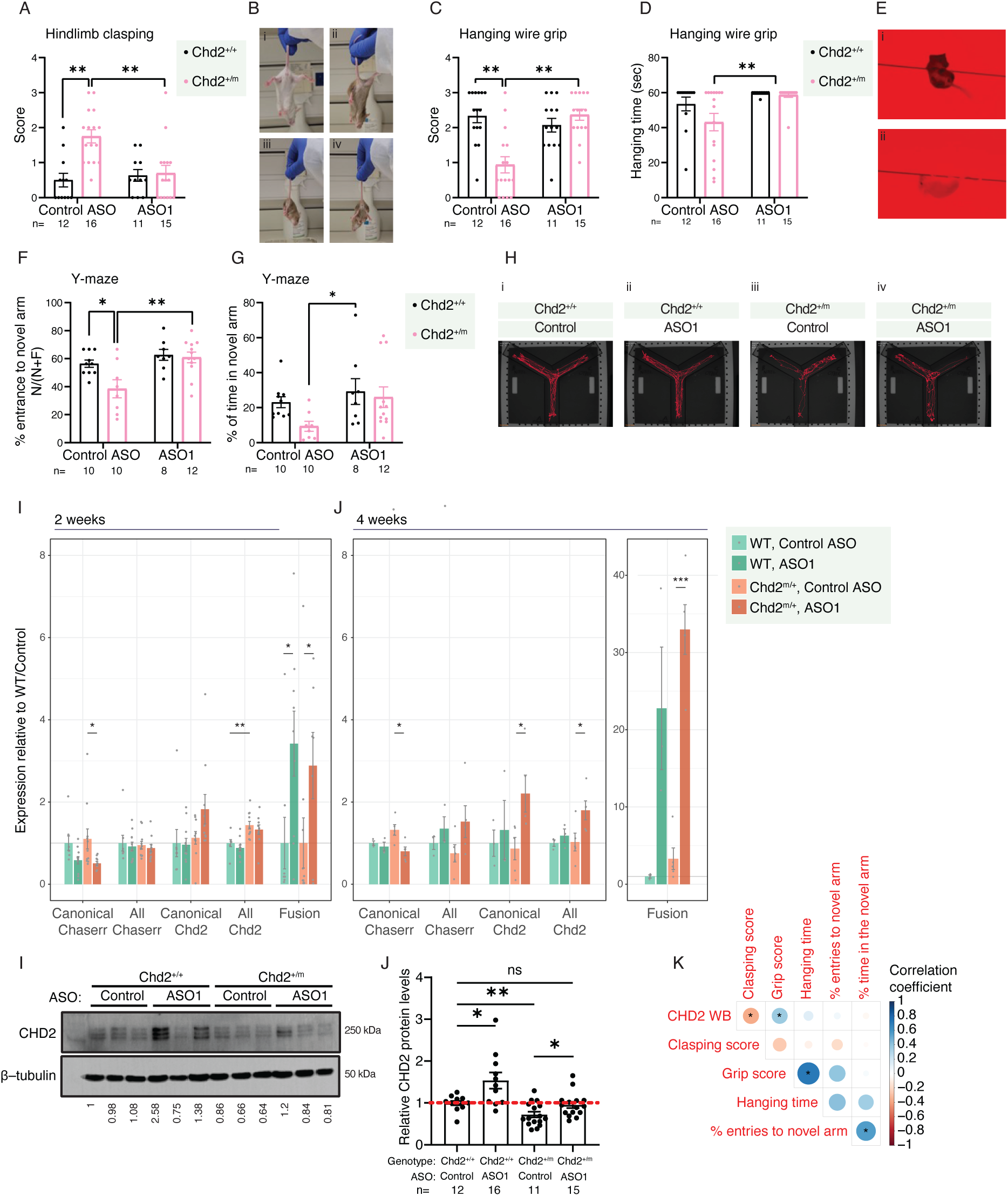
ASO1 rescues behavioral phenotypes in a model of CHD2 haploinsufficiency. **(A)** Scores of hindlimb clasping, ranging from 0 (normal, no clasping) to 3 (severe clasping, full hindlimb retraction) for mice with the indicated genotypes injected with the indicted ASO. P-values computed by two-way ANOVA. **(B)** A visual scale of the standardized assessment of clasping behavior in mice: i - a mouse exhibiting no clasping (score 0); ii - a mouse clasping one hindlimb (score of 1); iii - a mouse clasping both limbs (score of 2-3) (see Methods for the full details of the scoring theme). **(C-D)** Score (C) and time spent on the wire (D) for mice with the indicated genotypes injected with the indicated ASOs. **(E)** Examples of hanging wire behavior in mice; top: a mouse struggling to balance, receiving a score of 0–1 (see Methods); bottom: a mouse able to balance and traverse the wire, receiving a score or 2–3. **(F-G)** Fraction of entries to the novel arm of the maze (F) and fraction of time spent in the novel arm (G) for mice with the indicated genotype injected with the indicated ASO. **(H)** Examples of tracks of individual mice with the indicated genotype/ASO combinations. **(I-J)** Western blot (I) and quantification of a number of blots (J) for CHD2 levels in the brain of mice from the indicated genotypes injected with the indicated ASOs, and harvested at 6 weeks of age. **(K)** Pearson’s correlation coefficients and significance for the correspondence between CHD2 Western blot-derived levels (WB) and the indicated scores, among the Chd2_+/m_ mice injected with the control ASO or ASO1. In all panels, error bars indicate SEM; *- P<0.05, ** - P<0.005, *** - P<0.0005.

We also examined the molecular consequences of ASO1 introduction. At the ages of 2 and 4 weeks, we found that ASO1 increases fusion transcript formation, and at 4 weeks there was a significant increase in overall Chd2 mRNA levels throughout the brain (**Fig. 6I-J**). At these stages, we did not yet observe a significant increase in CHD2 protein levels (**Fig. S5B**), possibly because of a limited sample size and variability in CHD2 expression in whole-brain samples.

Levels of CHD2 protein were significantly increased in the brains of the mice subjected to the behavioral tests at the endpoint of the experiment (6 weeks, **Fig. 6I-J**). When we considered correlations between the performance of the *Chd2*^+/m^ mice in the three tests and CHD2 levels (**Fig. 6K**), we found that performance in the three tests was positively correlated with each other (note that WT mice have lower clasping scores and higher grip scores), with the correlations in performance in hanging wire Grip and Y-Maze tests being significant, and clasping and grip scores correlated with CHD2 protein levels.

### Fusion-induction potential is found in other tandem gene pairs

To evaluate if the principle of fusion formation from tandem genes, their regulation by MCR-related motifs, and their induction by ASOs extend beyond CHASERR and CHD2, we analyzed the RefSeq gene annotation resource. We first examined the prevalence of RRAUGG motifs.

Last exons of human protein-coding genes (see Methods) were slightly but significantly enriched for these motifs compared to randomly shuffled sequences of last exons (enrichment factor 1.1, P<10^−5^). When examining the position of the motifs the slight enrichment was found throughout the exon (**Fig. 7A**). When examining sequence conservation however, and comparing to shuffled versions of the RRAUGG motifs, preferential conservation (based on averaged PhyloP scores^11^) was mostly found in the first 100 bases of the last exon, with the motif being in the 96^th^ percentile of conservation, relative to 180 possible permutations, in the first 50 nt of the last exon, and 86^th^ percentile in positions 51–100 (**Fig. 7A**). The motif was also over-represented to a similar degree and preferentially enriched relative to its permutations in internal exons, and in first exons, suggesting that it does not carry out a preferential role in last exons, but rather more likely to play a general role in exon definition. Notably, the motif is contained within the TISU motif SAASATGGCGGC ^17^, which can explain its preferential enrichment in the first exons. Importantly, there is no evidence of translation initiation at the RRAUGG motifs in the last Chaserr exon (**Fig. S3C**).

**Figure 7.**
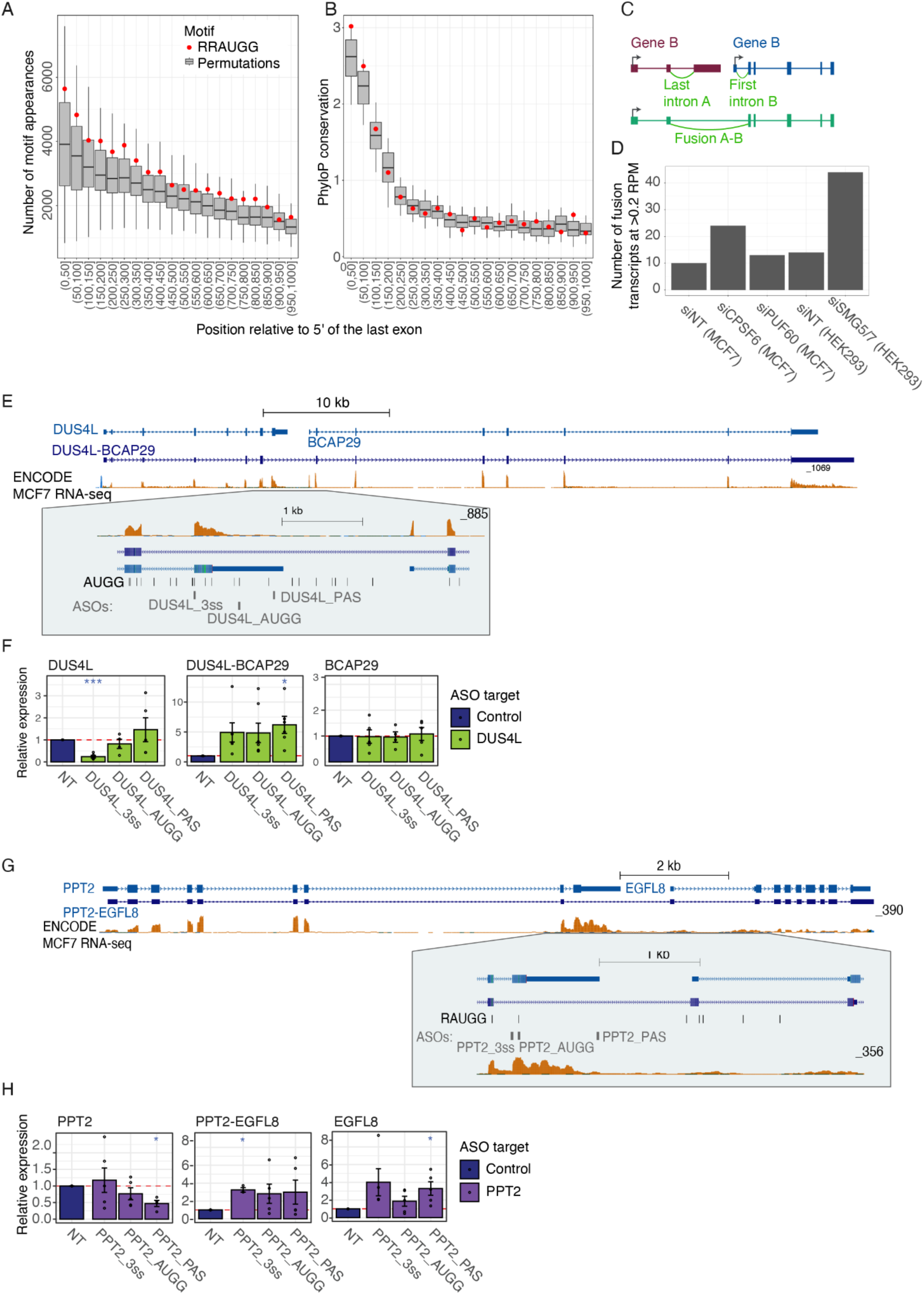
Fusion induction potential is found in other genes. **(A-B)** Number of appearances (A) and PhyloP score based on 100-way whole genome alignment of vertebrate genomes (B) of either the RRAUGG motif or one of its 180 permutations in the indicated position bin relative to the 3’ splice site of all the human protein-coding genes annotated in B. **(C)** Illustration of the scheme used to measure formation of fusion transcripts. **(D)** Number of fusion transcripts detected at the 0.2 RPM threshold in the indicated cells. MCF7 data is from this study, and HEK293 data are from^31^. **(E)** Overview of the DUS4L-BCAP29 locus with the MANE gene models and the fusion transcript annotated in GENCODE. RNA-seq read coverage on the ‘plus’ strand from the ENCODE project is shown. **(F)** qRT-PCR measurements of the indicated transcripts in MCF7 cells treated with the indicated ASOs. **(G)** As in A, for the PPT2-EGFL8 locus. **(H)** As in F for the PPT-EGFL8 gene pair.

To evaluate the extent of fusion transcript formation, we identified in the RefSeq annotations 2937 A-B gene pairs transcribed from the same strand where the intergenic region was 6 kb or less. We then mined RNA-seq datasets from our MCF7 cells with CPSF6 and PUF60 KD, as well as HEK293T cells following NMD inhibition. Specifically, we looked for reads supporting the splicing of the last intron of gene A, the first intron of gene B, or the potential fusion transcript (**Fig. 7B**). In the MCF7 control datasets there were 10 pairs with evidence for fusion expression at >0.2 reads-per-million-mapped-reads (RPM), while 14 pairs showed evidence in the HEK293 WT controls. Following KD of CPSF6 and PUF60, 26 and 31 pairs, respectively, were induced by >2-fold, and 24 and 13 fusion transcripts were expressed at >0.2 RPM, including CHASERR- CHD2 (**Fig. 7C**). We analyzed RNA-seq datasets from human cells following NMD inhibition mediated by SMG5 knockdown in SMG7 KO cells^31^ to investigate whether some of these fusions are occasionally produced endogenously and subset to NMD. Here, 76 gene pairs were induced by at least 3-fold compared to baseline control samples, with 44 reaching >0.2 RPM (**Fig. 7D**). We conclude that additional gene pairs have the potential to form fusion transcripts and that expression of these transcripts is limited by CPSF6, PUF60, and NMD.

We then selected four of the tandem gene pairs that were expressed in MCF7 cells and used ASOs to block their splice sites, PAS sequences, and/or selected AUGG sequences. The gene pairs were selected based on the relative proximity of the end of one to the start of another, evidence of transcription in the intergenic region, and AUGG motifs were selected based on their relative conservation. Of the 11 ASOs tested, two, targeting *DUS4L* and *PPT2* led to a statistically significant 6.2- and 3.2-fold increase in the formation of fusion transcripts *DUS4L*- *BCAP29* (annotated as ENST00000673970.1) and *PPT2*-*EGFL8* (annotated as ENST00000422437.5), respectively (**Fig. 7E-H**). As in the case of *Chaserr*-*Chd2*, the ASOs have uniformly increased fusion expression while having variable effects on the expression of the upstream gene (**Fig. 7F,H**). We conclude that ASOs targeting upstream genes in tandem gene pairs can be used for inducing the formation of fusion transcripts between gene pairs, which can result in the induction of overall expression of the downstream gene.

## Discussion

The current estimate is that there are >500 haploinsufficient genes in the human genome^3^. Haploinsufficiency is particularly common in neurological diseases with causative genes involved in transcription and/or chromatin biology, with >65 different chromatin-associated genes being haploinsufficient ^4^. Gene therapy for such disorders often faces challenges such as delivery efficiency, high costs, and safety concerns ^32^, and indeed, to the best of our knowledge, none are currently in the pipeline for most genetic conditions, including *CHD2*. Expression levels are difficult to control in the context of gene therapy, which is a major issue in the case of *CHD2* where over-production appears to have more adverse consequences than haploinsufficiency^9,33^. Unlike gene editing, which is irreversible, RNA-based approaches allow for controlled regulation of gene expression, providing a reversible and targeted method to increase protein levels where needed. ASOs, rendered very stable by chemical modifications, are a successful and clinically proven approach for gene regulation, with clinically approved ASO-based treatments given every 4-6 months^6^. Despite their evident success, traditional ASOs are limited in the types of gene regulation they can confer, constraining the implementation of this effective approach in a broader range of health conditions. We describe here a novel approach that circumvents some of the limitations of ASOs and facilitates the boosting of gene expression for genes that have a close upstream tandem gene. As we show, targeting the last exon of the upstream gene can lead to the formation of fusion transcripts, and those transcripts can facilitate the production of additional protein copies, which is particularly relevant for haploinsufficient genes.

We show that targeting different regions within and in proximity to the last exon of *Chaserr*, as well as that of two other genes, is sufficient for induction of the fusion transcript. While it is less surprising that inhibiting *Chaserr* polyadenylation or the splicing of its last exon in the context of pre-existing transcription downstream of it promotes fusion formation, it is more surprising that ASO3 which targets regions of the *Chaserr* MCR located >160 nt away from the last 3’ splice site and >125 nt from the CPA site is at least as effective as ASOs targeting directly either the 3’ splice site of the PolyA signals (**Fig. 4E**). We suggest that the *Chaserr* MCR acts as coordination region, promoting both splicing and polyadenylation, and so the blockage of RRAUGG motifs is more efficient in exclusion of the last exon than separately targeting splicing or polyadenylation. It is possible that this region, which is highly conserved in evolution (**Fig. 1A**) is optimized to perform this function, and so its disruption results in more efficient fusion transcript formation than what we can obtain for other gene pairs, where ASOs targeting 3’ splice sites, polyA signals, or AUGG motifs lead to a more modest increases in fusion transcripts (**Fig. 7**). It is also possible that MCR-targeting ASOs are more efficient in binding to CHASERR transcript than other ASOs due to limitations imposed by sequence composition.

The *Chaserr* MCR contains several RRAUGG motifs, present in the last *Chaserr* exons of homologs throughout vertebrates^10^, suggesting it has a conserved role. We found evidence for formation of *chaserr*-*chd2* fusion during the later stages of embryonic development in zebrafish (4–8 hpf, starting from the dome stage, **Fig. S6A-B**, based on data from^34^) and in the zebrafish brain (based RNA-seq data from^35^). Interestingly, the ratios between the expression levels of *Chaserr*, *Chd2* and the fusion transcripts are overall similar in all the different systems we studied in human, mouse and zebrafish, with *Chaserr* expression ∼2-10-times higher than that of *Chd2*, and ∼100 fold higher than the fusion (**Fig. 1E**, **2C, S7B**), suggesting that in control conditions ∼1% of the *Chaserr* transcription events result in fusion transcript formation. Upon conditions such as neuronal activation, an increase in fusion formation can thus produce a number of fusion mRNA molecules comparable to those produced by the canonical *Chd2* mRNA, and the differential translation control offered by the fusion-derived vs. the canonical 5’ UTR can then further tune CHD2 protein levels. Importantly, endogenous functions of the fusion transcript can explain the conservation of the sequences in the first four exons of *Chaserr*, which is a 5-exon transcript in various vertebrate species^7^, as these exons form the 5’UTR in the fusion transcript and are likely important for translational control of the CHD2 protein produced from it.

The RRUAGG motifs in the *Chaserr* MCR events are presumably recognized by protein factors. We previously used mass spectrometry and a pulldown of an *in vitro* transcribed last *Chaserr* exon and identified proteins that bind these motifs, yet knockdown of key hits from that screen, *Dhx36*, *Rbm3*, and *Zfr*, did not increase fusion formation. It is possible that the motifs are recognized in a specific endogenous context and/or are associated with redundant regulatory factors.

Several factors currently limit the widespread application of our approach. One limitation is that we are up-regulating both the WT and the mutant copy of *Chd2*. In our mouse model the *Chd2*^m^ allele-derived mRNA is overall not preferentially destabilized compared to the WT copy^30^, and similar observations have been made in iPSC-derived cells from human CHD2 patients^36^. While the available evidence does not suggest that increased production of the mutated allele is consequential in the case of CHD2, with haploinsufficiency being the most likely mode of pathogenecity^37^, it can not be ruled out as a point of concern at this point.

Another limitation is that in most cases, if the upstream gene is expressed well enough to provide a relevant reservoir of additional transcripts, it is typically a protein-coding gene, and those have ∼8 exons on average, and thus the 5’ UTR preceding the downstream ORF in the fusion transcript will permit limited downstream ORF translation according to the currently accepted model of translation re-initiation. IRES-dependent translation, the rules of which are poorly understood in mammalian mRNAs, can be relevant for translation in fusion mRNAs if it is supported by downstream exons in multi-exon mRNAs. As we show here, the 118 aa ORF in *Chaserr* still allows efficient and therapeutically relevant downstream translation. While in a reporter setting we see that the *Chaserr*-based 5’ UTR supports translation that is >20-fold less efficient than one supported by the Chd2 5’ UTR, the significant increases in CHD2 protein levels upon ASO1 administration, without consistent changes to the levels of the canonical Chd2 mRNA, and the evidence that the fusion mRNA is enriched at polysomes both in our experiments and in the public data (**Fig. 3G** and **Fig. 3H**) both suggest that the endogenous fusion transcript can be efficiently translated. Ribo-seq, typically a method of choice for measuring translation efficiency, is not applicable for measuring the translation efficiency of the fusion transcript, as the coding sequence of the fusion is identical to that of the canonical Chd2. Notably, the Chaserr uORF begins from an AUG that is very close to the first nucleotide of the Chaserr RNA, which some transcript isoforms apparently do not contain (**Fig. S3C**). The AUG codon is found in a strong TISU context, which is highly enriched in very short 5’ UTRs^17,18^.

Notably, the translation of TISU elements is also highly regulated ^38,39^. Therefore, there is ample room for regulation of the translation of the ORF embedded in *Chaserr* and in the fusion transcript, which in turn can regulate translation of the CHD2 protein from the fusion transcript in ways that can be elaborated in future studies.

As we show, the *Chaserr*-*Chd2* fusion transcript that is induced by ASO treatment is also formed, albeit at low levels, in untreated human and mouse cells. Neuronal activation in culture increases the formation of this fusion. We further show that in a gain-of-function model where the fusion transcript is formed constitutively, there are changes in gene expression and chromatin accessibility that become apparent in activated primary neurons. Surprisingly, we find significantly increased chromatin accessibility, possibly due to increased CHD2 protein levels and enhanced chromatin remodeling in peaks associated with genes that are downregulated on the transcriptional level in activated neurons. It is possible that the enhanced accessibility reflects changes in nucleosome positioning that affect robust transcriptional induction upon activation. A clearer view will emerge once the genomic occupancy of CHD2 can be mapped in these cells, as our attempts to use Cut&Run to profile CHD2 occupancy have so far not been successful. Pinpointing the specific functional contribution of the fusion transcript is challenging as methods that disrupt fusion transcript formation are likely to also disrupt nascent transcription through the *Chaserr*/*Chd2* locus. Indeed the GapmeRs we use to target the intergenic region between Chaserr and Chd2 reduce the levels of the fusion transcript, but also the canonical *Chaserr* transcripts (**Fig. S1B**).

Our results suggest that CHASERR can regulate CHD2 through several distinct mechanisms (**Fig. 8**). First, the promoters of CHASERR and CHD2 both contact a series of shared enhancers found in the gene desert upstream of CHASERR, where several enhancers regulating CHD2 are found, and some have been recently validated in human cells^33,40^. We have previously shown that deletion of the *Chaserr* promoter increases contact frequency between these enhancers and the *Chd2* promoter, suggesting promoter competition as one possible mechanism of regulation. Second, it is possible that transcription through the *Chaserr* locus, which, as we show here, continues at least to the promoter of *Chd2*, represses the *Chd2* promoter through mechanisms related to those of other interfering transcripts, mostly characterized in yeast^41^. This mechanism can explain why deletion of the *Chaserr* gene body, or knockdown of its expression with GapmeRs both lead to increase in *Chd2* levels^7^, which is unlikely to be explained by promoter competition. One of these mechanisms, or another, presently unknown, leads to an over-production of CHD2 in human patients that carry heterozygous deletions in the first three exons of CHASERR, which results in CHD2 up- regulation, and a severe neurodevelopmental disorder^9^, mirroring the severe phenotypes of *Chaserr*^+/–^ mice. The same mechanisms likely underpin the embryonic lethality of *Chaserr*^−/−^ mice^7^. Crucially, fusion-transcript–related mechanisms cannot explain these phenotypes, as *Chaserr* promoter removal results in complete loss of *Chaserr* expression, which in turn prevents production of a *Chaserr*-*Chd2* fusion transcript. Indeed, we found no fusion reads in RNA-seq data obtained from *Chaserr*^+/–^ mice^7^ . Fusion formation with the *Chd2* mRNA thus constitutes a separate mechanism through which *Chaserr* can regulate *Chd2*, and its perturbation is associated with milder phenotypic outcomes, as the MCR-S and MCR-L mouse models are viable and fertile, with only a mild reduction in number of homozygous *Chaserr*^MCR-S/MCR-S^ pups compared to the one expected by chance (**Fig. S2B**).

**Figure 8.**
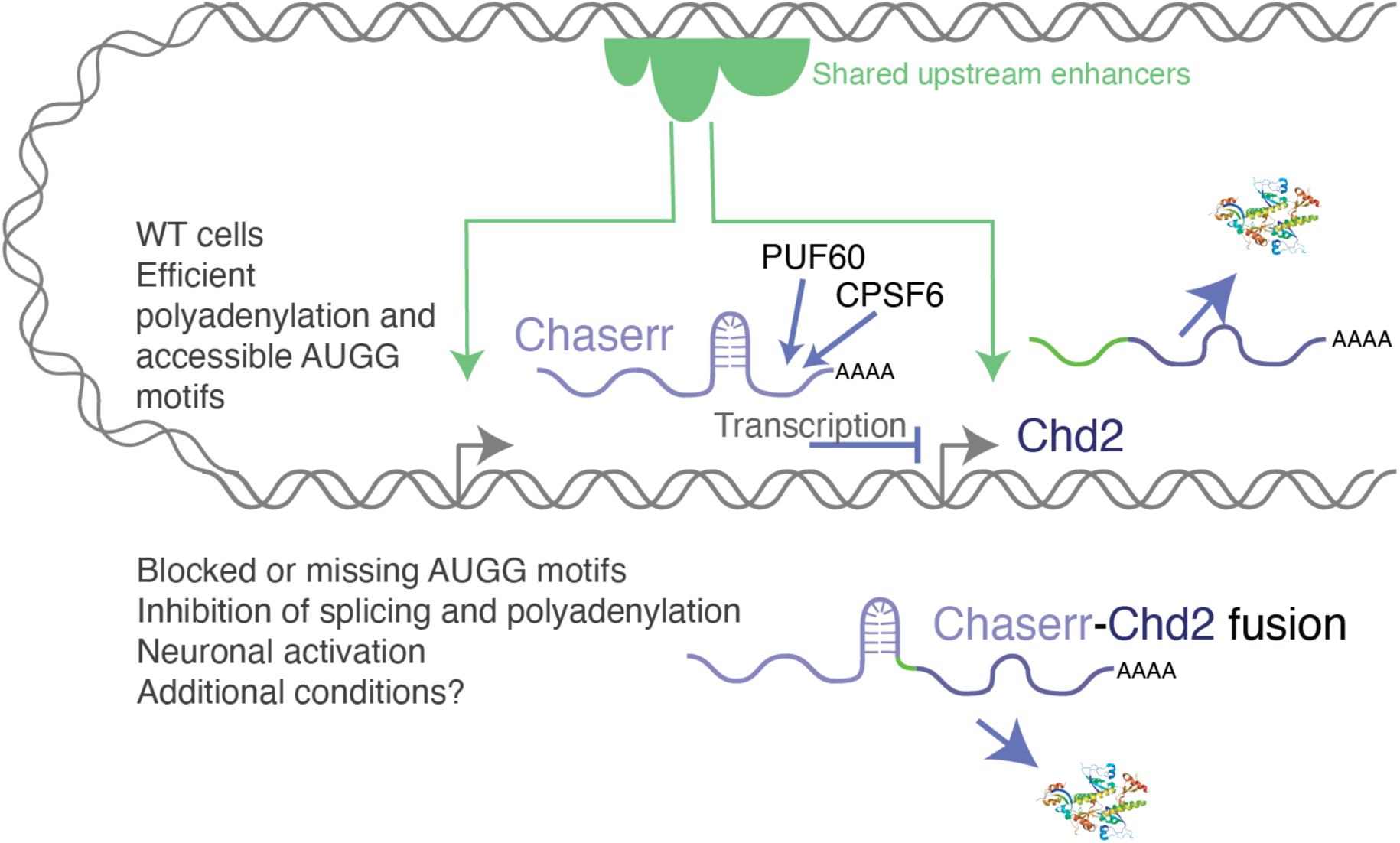
Model for multi-layer regulation of CHD2 by CHASERR.

CHD2 haploinsufficiency affects >440 known individuals (https://www.curechd2.org/map), with the vast majority of cases being young individuals diagnosed recently via exome sequencing, suggesting the true prevalence of the condition is significantly higher. Targeting *CHASERR* as a repressor of *CHD2* is currently considered a major direction for developing therapeutics for this condition, which currently has no other specific therapeutics ^42^. A major question regarding CHD2 haploinsufficiency is whether CHD2 levels mostly affect brain development or functions of the postnatal brain. We show here three robust behavioral phenotypes exhibited by the *Chd2*^+/m,^ which are significantly rescued by administration of ASO1, suggesting both that we have a model that can be used for measuring CHD2 activity and that affecting CHD2 levels in the postnatal brain is sufficient for rescuing specific manifestations. The mild phenotype of *Chaserr*^MCR^ models further suggests that induction of the fusion transcripts in *CHD2*^+/–^ is unlikely to lead to the strong adverse effects of CHD2 over-production observed in *CHASERR*^+/–^ individuals.

## Materials and Methods

List of reagents used in this paper can be found in **Table S1**.

### Tissue culture and transfections

NIH/3T3 and MCF-7 cell lines (obtained from ATCC) were routinely cultured in DMEM containing 10% fetal bovine serum and 100 U penicillin/0.1 mg ml−1 streptomycin at 37 °C in a humidified incubator with 5% CO2. Cell lines were routinely tested for mycoplasma contamination and were not authenticated. All ASOs (Integrated DNA Technologies, **Table S2**), LNA GapmeRs (Qiagen, **Table S2**) and siRNAs (Dharmacon SMARTpool, **Table S3**) were transfected into NIH/3T3 and MCF-7 cells using DharmaFECT 1 and DharmaFECT 4 respectively. Transfections were carried out according to the manufacturer’s protocol, to a final concentration of 25 nM. Luciferase plasmid transfections were performed using PolyEthylene Imine (PEI) (PEI linear, M*r* 25,000, Polyscience), to a final concentration of 100 ng in NIH/3T3 cells. Endpoints for all experiments were at 48 hr post transfection. All ASOs and siRNAs used in this study are listed in Tables S2-S3.

### NMD Inhibition

NIH/3T3 and MCF-7 were transfected with either ASO1 or a non-targeting control antisense oligonucleotide (ASO) at a final concentration of 25 nM using DharmaFECT reagents according to the manufacturer’s instructions. After 48 hours, cells were treated with either 200 μg/mL cycloheximide (CHX C7698; Sigma-Aldrich) to inhibit nonsense-mediated decay (NMD) or an equivalent volume of ethanol (EtOH) as a vehicle control for 8 hours. Following treatment, total RNA was extracted and expression levels of *Chaserr*, *Chd2*, and the *Chaserr*-*Chd2* fusion transcripts were measured by RT-qPCR and normalized to the housekeeping gene actin using the ΔΔCt method.

### Subcellular fractionations

Fractionations were performed as previously described ^43^. Briefly, cells were collected and lysed in buffer RLN (50mM Tris•Cl pH8, 140mM NaCl, 1.5mM MgCl_2_, 0.5% v/v NP-40, 10mM EDTA, 1mM DTT, 10U/ml RNase Inhibitor), The supernatant was collected as the cytoplasmic fraction and the nuclear pellet was washed in Buffer S1 (250mM Sucrose, 10mM MgCl_2_, 10U/ml RNase Inhibitor) and layered on top of a cushion of buffer S3 (880mM Sucrose, 0.5mM MgCl_2_, 10U/ml RNase Inhibitor). RNA was extracted from the different fractions using BioTri reagent (Bio-Lab 959758027100).

### Polysome profiling

NIH/3T3 were transfected with ASO1 at a final concentration of 25nM. After 48 hours cells prepared for polysome profiling as previously described ^44^. Briefly, cells were incubated in 100µg/ml CHX for 10 minutes at 37°C, and subsequently washed three times with ice-cold Phosphate Buffered Saline (PBS) supplemented with 100µg/ml CHX. Cells were collected by scraping and centrifugation for 5 minutes at 500 x g at 4°C. Pellets were resuspended and lysed for 10 minutes on ice in 1ml polysome extraction buffer (PEB) (20nM Tris-HCl pH 7.5, 100mM KCl, 5mM MgCl2, 0.5% Nonidet P-40, 100µg/ml CHX, 10U/ml RNase Inhibitor, 1x protease inhibitor). Lysates were cleared by centrifugation and a total of 750µg RNA was layered to the top of a 10-50% sucrose gradient (prepared using the BioComp Gradient Master) and centrifuged for 90 minutes in a SW41Ti swinging bucket rotor at 39,000 rpm at 4°C. Gradients were fractionated on the Triax FlowUV/Fluorescence Gradient Profiling system by collecting 1ml/min fractions into 24 tubes. RNA was extracted from each gradient fraction using BioTri (Bio-Lab 959758027100), and the distribution of RNA for each gene of interest was determined by RT-qPCR.

### Luciferase assays

Reporter plasmids were constructed by cloning the alternative 5’UTR sequences in a psiCheck2 plasmid. Luciferase activity was measured 48 hr post transfection using the Dual-Glo Luciferase Assay System (Promega). Following cell lysis, 20µl of extract was placed in a black 96 well plate and activity of Firefly and Renilla luciferase was measured using the GloMax® Explorer Multimode Microplate Reader from Promega. A relative response ratio, from RnLuc signal/ FFLuc signal, was calculated for each sample.

### Animals

The study was conducted in accordance with the guidelines of the Weizmann Institutional Animal Care and Use Committee (IACUC). All mouse strains were bred and maintained at the Veterinary Resources Department of the Weizmann Institute. Adult mice, aged 6 to 8 weeks, were used in the study.

### Transgenic mice generation

The CRISPR KO mice were generated as in ^45^. Mice were generated by standard procedures at the Weizmann transgenic core facility using two single guide RNAs (sgRNAs, **Table S4**) with recognition sites (chr7:73,544 and chr7:73,544,263 mm10 assembly). We generated two deletions using these guides: MCR-S: chr7:73,544,266-73,544,373 and MCR-L: chr7:73,544,257-73,544,416. Injections were performed into pure C57BL/6 mice. Mice were identified by genotyping and sequencing. Lines were bred and maintained on C57BL/6 background at the Veterinary Resources facility of the Weizmann Institute. *Chd2*^m/+^ were generated and described previously, originally on a mixed background ^7^, and then backcrossed to the 129 background, which revealed behavioral phenotypes ^30^. Mice crossed to the 129 background for at least 10 generations were used in this study.

### RNA sequencing

PolyA selected libraries were generated using the CORALL mRNA-seq V2 kit (Lexogen 177.96) according to the manufacturer’s protocol. The samples were sequenced using the Illumina NovaSeq X instruments. RNA-seq reads were mapped to the mm10 or hg38 genome assemblies using STAR ^46^ which generated read coverage bigWig files, splice-junction–specific counts, and gene read count files, which were normalized to reads-per-million (RPM) in R. Differential expression was computed using DeSeq2 ^27^. Visualization of read coverage and intron-spanning reads was with UCSC Genome Browser ^47^ and Integrative Genome Viewer ^48^. Sashimi plots for splicing were visualized using sashimi ^49^.

### Long read RNA-seq data analysis

Long read RNA-seq data generated by Oxford Nanopore sequencing was downloaded from the ENA database project PRJEB44348, and mapped to the hg38 human genome assembly using minimap2 ^50^. BAM files were converted to BED12 representation using BEDtools ^51^ bamtobed and reads overlapping the intergenic region between CHASERR and CHD2 were inspected using IGV ^48^.

### RT-qPCR

RNA was extracted using BioTri (Bio-Lab 959758027100) and cDNA prepared using the qScript flex cDNA Synthesis Kit (Quantabio 95049) according to the manufacturer’s protocol.

Expression levels were measured on the ViiA 7 Real-Time PCR System (Applied Biosystems) using Fast SYBR™ Green Master Mix (Thermo 4385614). Results are displayed as relative expression levels determined by the ΔΔCt method. Primer sequences are available in **Table S5**.

### 5’ RACE

RNA from WT or MCR homozygous mice was used for cDNA generation with gene specific primers targeting either the second exon of Chd2 or the last exon of Chaserr using SMARTScribe reverse transcriptase (Takara 639537) in the presence of a template switch oligo. The cDNA was amplified using primers targeting the Illumina R1 and R2 regions adding barcodes and the p5 and p7 regions to the amplicon. Libraries were sequenced using the Illumina NovaSeq X instrument. Primer sequences are available in **Table S5**.

### Protein extraction and Western Blot

Proteins were extracted using RIPA Buffer (150mM NaCl, 1% NP-40, 0.5% Deoxycholate, 0.1% SDS, 50mM TRIS pH 8, 1mM DTT). Protein expression was analysed by SDS-PAGE followed by Western blot. The complete list of primary and secondary antibodies and relative dilutions is available in **Table S1**.

### Cleavage Under Targets and Release Using Nuclease (CUT&RUN)

CUT&RUN reactions were performed following the V3 of the protocol ^52^, with minor modifications. An equal number of cells (up to 350,000 for each sample) was harvested using Trypsin (0.05%) solution and washed once with 1X PBS at room temperature. Cells were then washed at three times at RT in Wash Buffer (20mM HEPES-NaOH pH 7.5, 150mM NaCl, 0.5mM Spermidine supplemented with EDTA-free Protease Inhibitor Cocktail (APExBIO)).

Following the third wash, cells were resuspended in 1 mL Wash Buffer and incubated while gently rotating with 20 µL of Concanavalin A-coated magnetic beads (EpiCypher), which had been previously washed, activated and resuspended with Binding Buffer (20mM HEPES-NaOH pH 7.5, 10mM KCl, 1mM CaCl_2_, 1mM MnCl_2_). After 10 minutes incubation, the buffer was removed, and the beads were resuspended in 150 µL of Antibody Buffer (Wash Buffer with 0.1% Digitonin (Sigma) and 2mM EDTA pH 8) containing the proper dilution of each antibody of interest: Total Pol2 (Rpb1 CTD (4H8) Mouse mAb #2629), Ser2P and Ser5P ^53^ (a kind gift from Hiroshi Kimura) . The cells were then incubated with each antibody, with gentle rotation overnight with a 180° angle at 4°C. On the next day, cells were washed twice with ice-cold Dig-Wash Buffer (Wash Buffer with 0.1% Digitonin), resuspended in 150 µL of Dig-Wash Buffer supplemented with 1 µL of custom-made pA/G-MNase every mL and left rotating for 1h at 4°C.

After that, cells were washed twice with Dig-Wash Buffer, resuspended in 100 µL of Dig-Wash Buffer, and placed into a thermoblock sitting on ice. To initiate the cleavage reaction, 2 µL of a freshly diluted (from a 1M stock) solution of 100mM CaCl_2_ was added to each tube, and the tubes were left at 0°C for 30 minutes. To halt the reaction, 100 µL of 2X STOP Buffer (340mM NaCl, 20mM EDTA pH 8, 4mM EGTA pH 8, 0.05% Digitonin, 100 µg/mL RNAse A, 50 µg/mL Glycogen) were added to each tube, and cleaved DNA fragments were released by incubating the samples for 30’ at 37°C. The tubes were then centrifuged for 5 minutes at 4°C / 16,000g and the supernatant was collected. Finally, the DNA was purified by standard Phenol/Chloroform extraction using the 5PRIME Phase Lock Gel Heavy tubes (QuantaBio), and the success of the CUT&RUN reaction was assessed by running the positive control (H3K27me3 or H3K4me3) on a Tapestation (Agilent Technologies) using a High Sensitivity D1000 ScreenTape (Agilent Technologies).

### HCR

All probes were designed and purchased from Molecular Instruments (**Table S6**) and the RNA-FISH protocol was carried out according to Molecular Instrument’s protocol for Mammalian Cells on a Chambered Slide. Briefly, NIH/3T3 cells were grown in chambered slides coated with 0.01% (w/v) poly-D-lysine for 24 hours before transfection with ASO1 to a final concentration of 25nM. 48 hours post-transfection cells were fixed in 4% PFA at room temperature (RT) for 10 minutes and rinsed twice in PBS. Cells were then permeabilized in 0.1% Triton at RT for 30 minutes, washed twice in PBS for 3 minutes and incubated in 70% EtOH at -20°C overnight. On the next day, cells were blocked with Molecular Instruments AB buffer for 1 hour at RT with shaking, followed by three washes in PBS for 5 minutes at RT and rinsed twice with 5x SSCT. Cells were first pre-hybridized for 30 minutes at 37°C in HCR hybridization buffer with 100 ug/ml salmon sperm DNA (SSD). Probes that targeted the first four exons of Chaserr and the full Chd2 mRNA were then added in hybridization buffer containing 100 µg/ml SSD, and hybridization was carried out overnight at 37°C. On the next day, cells were washed four times in HCR wash buffer for 5 minutes at 37°C, after which the HCR Amplification Buffer containing DNA hairpins and SSD was applied for 90 minutes at RT to achieve linear amplification. Cells were then washed five times in 5x SSCT for 5 minutes at RT and nuclear staining was performed with DAPI (Thermo Fisher Scientific). Fluorescent imaging was carried out using a Zeiss Spinning Disk confocal microscope.

### Primary neuronal culture

Embryonic cortices (embryonic day 16.5–17) were dissected from 5–6 embryos of mixed sex and pooled for a single replicate culture. Trypsin (0.25%) was used to dissociate tissue through a 10-minute incubation at 37°C. Digestion was terminated with the addition of ovomucoid (trypsin inhibitor, Worthington). Digested tissue was gently triturated through a P1000 pipette to release cells and passed through a 40 µm filter. After trituration, the cell suspension was transferred to a 15 mL Falcon tube and centrifuged for 5 minutes at RT / 154×g. After centrifugation, the cloudy supernatant was carefully removed with a pipette and discarded. The pellet was resuspended in 10 mL DMEM medium and added to uncoated 90 mm Petri dishes. The plates were incubated at 37°C, 5% CO2 humidified incubator for 30 minutes. During this incubation, glial cells and other unwanted debris adhere to the bottom of the plate while cortical cells are retained in the media and recover from the trituration. The supernatant was carefully removed without disturbing attached glial cells and unwanted cell debris at the bottom of the plate and moved into a new 15 mL Falcon tube using a pipette. The collected supernatant was then centrifuged for 5 minutes at RT / 154×g. Neurons were plated onto cell culture dishes pre-coated overnight with poly-L-lysine (20 µg/mL) and laminin (4 µg/mL) and were grown in neurobasal medium (Gibco) containing B27 supplement (2%), 1% PenStrep solution and glutaMAX (1 mM). Neurons were grown in incubators maintained at 37°C with a CO2 concentration of 5%. Cortical neuron culture was treated with 1 μM Tetrodotoxin (TTX) and 100 μM D-AP5 overnight, followed by 1 hr or 6 hr incubation with 55 mM KCl.

### ATAC-seq library preparation and analysis

ATAC-seq was performed as described ^54^. In brief, 25,000 cortical neurons were extracted in 25 µl of ice-cold cell lysis buffer (10mM Tris-HCl pH 7.4, 10mM NaCl, 3mM MgCl2, 0.1% IGEPAL CA-630). Tagmentation was performed using 2 μl of the Nextera Tn5 enzyme (TDE1). The libraries were sequenced with paired-end sequencing on NovaSeqX instruments.

ATAC-seq reads were mapped to the mm10 mouse genome assembly with Bowtie2 ^55^ and reads were extended and peaks called using MACS2 ^56^. Only libraries showing clear enrichment of accessibility around promoters were carried forward (n=34/40 samples of of three libraries from WT cells and five of *Chd2*^MCR-S/MCR-S^ ones, at least three libraries per condition). Peaks were called by combining all the libraries together using MACS2^56^, and the read counts in each peak were counted using Homer^57^. Differential accessibility was computed using DeSeq2 ^27^ which was executed through in Homer^57^.

### Analysis of RRAUGG motif representation

For analysis of RRAUGG counts, the last exons of all the human protein-coding genes were extracted and the appearance of RRAUGG motifs or each possible permutation was counted. For each appearance, the average PhyloP score was computed using bigWig files obtained from the hg38 assembly in the UCSC genome browser. Appearances were then gathered for each 50nt bin relative to the 3’ splice. A similar analysis was done for internal and first exons.

### Brain Tissue Preparation and Immunofluorescence

Brains were fresh-frozen in Tissue-Tek O.C.T. compound (Sakura, 4583) and sectioned at a thickness of 10 μm using a Leica CM3050 cryostat. Sections were fixed in 4% paraformaldehyde and subsequently subjected to blocking and permeabilization in a solution containing 5% donkey serum, 2% bovine serum albumin (BSA), and 0.1% Triton X-100 in phosphate-buffered saline (PBS). Primary antibody incubation was performed in the same permeabilization buffer using an anti-Chd2 (NBP2-92115-0.1ml, NUVOS) primary antibody overnight. Following washes, sections were incubated with a secondary antibody, Goat anti-Rabbit Alexa Fluor 647 (Abcam, ab150079). Nuclear staining was performed using DAPI (Thermo Fisher Scientific). Fluorescent imaging was carried out using a Leica DM4000 B microscope equipped with a Leica DFC365 FX CCD camera and Leica Application Suite (LAS) X software.

### Intracerebroventricular (ICV) Injection in Neonatal Mice

ICV injections were performed on postnatal day 2 (P2) mouse pups. Pups were immobilized by gentle restraint on a soft surface positioned above an ice-filled container and gently restrained using two fingers. A 33-gauge beveled needle (0–20 mm adjustable length, point style 4) was attached to a 10-μl Neuros Hamilton syringe (Model 1701, Gastight, Small RN, PN: 65460-06) was used to perform the injection. Injection coordinates were approximately 1 mm lateral to the bregma, 2 mm anterior, and 2 mm ventral. A total of 5 μl, containing 20 μg of ASO, was slowly injected into the right lateral cerebral ventricle. After injection, pups were promptly returned to the nest and monitored daily for survival and signs of stress.

### Behavioral tests

All behavioral tests were conducted between P28 and P36, and the mice were sacrificed at six weeks of age.

### Hindlimb clasping test

Hindlimb clasping was used as a behavioral assay to evaluate motor abnormalities and disease progression in mice. Mice were gently held upside down by the base of the tail and videotaped for 45 seconds. A severity score ranging from 0 to 3 was used to quantify hindlimb clasping behavior, with half-point scores granted for behaviors falling between defined severity levels.

Scoring criteria were as follows: 0 = hindlimbs splayed outward and away from the abdomen (normal); 1 = one hindlimb retracted inward toward the abdomen for ≥50% of the observation period; 2 = both hindlimbs partially retracted inward toward the abdomen for ≥50% of the observation period; 3 = both hindlimbs fully retracted inward toward the abdomen for ≥50% of the observation period. Higher scores were indicative of increased motor impairment.

### Hanging wire grip test

To assess neuromuscular function, mice were subjected to the hanging wire grip test. Each mouse was placed on a horizontal wire by its forelimbs and allowed to grasp the wire with its hindlimbs to stabilize itself. The duration for which each mouse could maintain its grip before falling was recorded, with a maximum cutoff time of 60 seconds. Longer grip durations were interpreted as indicative of better neuromuscular function. In addition to recording latency to fall, mice were assigned a performance score ranging from 0 to 3, with half-point increments granted for intermediate behaviors. Scoring was defined as follows: 0 = the mouse fell from the wire and was unable to stabilize itself; 1 = the mouse remained on the wire but exhibited unstable behavior; 2 = the mouse maintained a stable grip and did not fall; 3 = the mouse was stable and able to traverse the wire freely. Scores between 2 to 3 were considered within the range of normal behavior.

### Y-Maze Test

The Y-maze test was employed to assess spatial working memory and cognitive flexibility, commonly impaired in models of neurological disorders. The apparatus consisted of three arms (designated A, B, and C) of equal length and separated by 120° angles. During the training phase, each mouse was placed at the end of one designated arm and allowed to explore only two of the arms (start and familiar arm) for 15 minutes, while access to the third (novel) arm was blocked. Following a rest period of at least one hour, the mouse was returned to the maze with all three arms accessible for a 5-minute test session. Mouse behavior was recorded, and arm entries as well as the time spent in each arm were analyzed using an automated tracking system (Ethovision XT, Noldus, the Netherlands). A preference for the novel arm was interpreted as evidence of intact spatial memory and exploratory behavior.

### Data availability

All RNA-seq, ATAC-seq and Cut&Run data have been deposited to the GEO database accession GSE298095, reviewer token mhofqgwibhuprid)

## Acknowledgements

This study was funded by the European Research Council grant lncIMPACT and the Israeli Science Foundation grant 743/24 to I.U.

---

**Figure S1.**
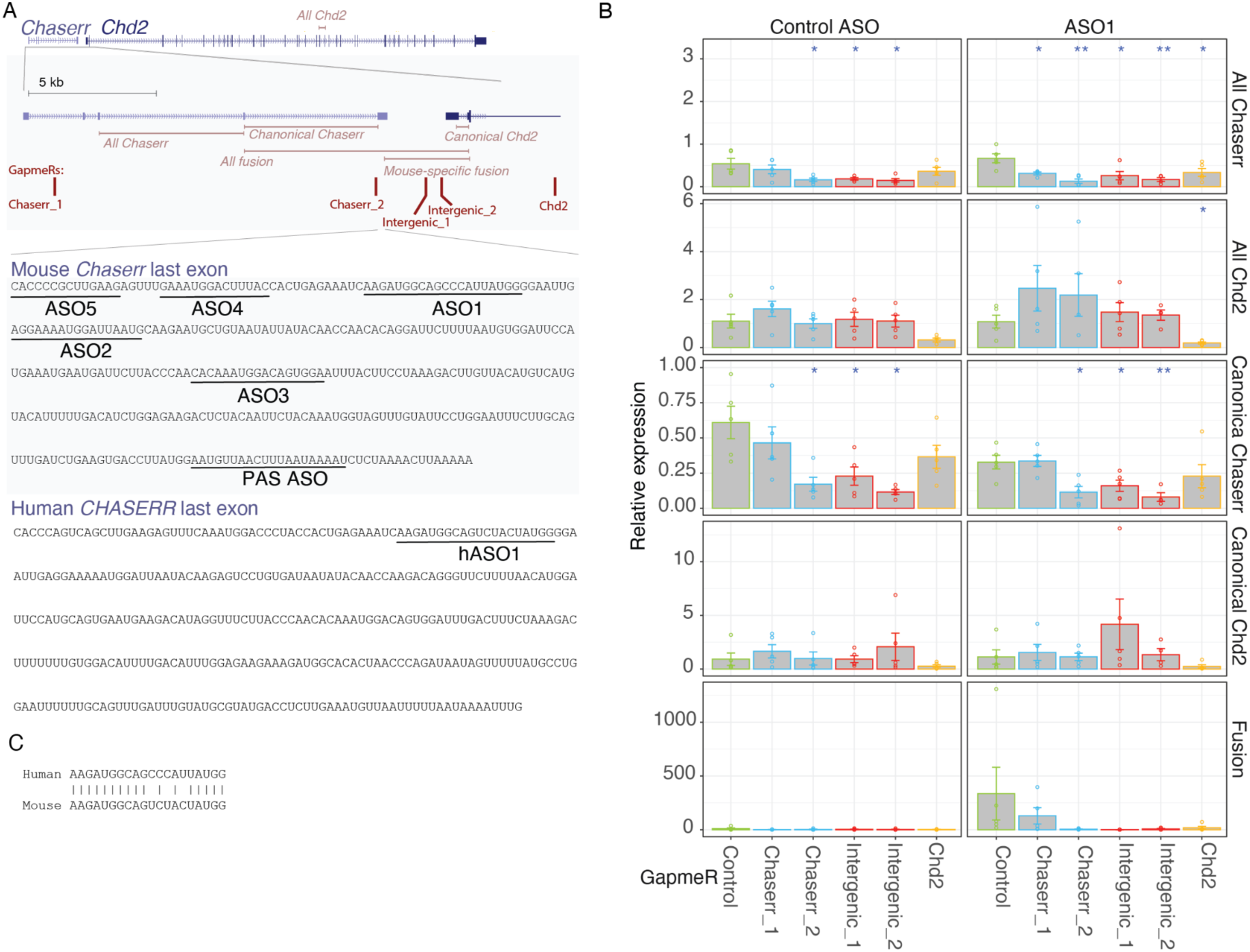
related to Figure 1. ASOs and GapmeRs targeting Chaserr and the intergenic region. **(A)** Positions of the different sequences within the Chaserr locus. Top: primers used throughout the manuscript for qRT-PCR in mouse cells and positions of the GapmeRs used. Bottom: sequences of the last exons of mouse and human *Chaserr*, respectively, marking the positions of the sequences targeted by the different ASOs used in this manuscript. **(B)** qRT-PCR measurements of the transcripts indicated on the right in cells co-transfected with the ASO indicated on top and GapmeRs indicated on bottom (positions of the sequences targeted by the GapmeRs are marked in A). Each transcript is normalizedto untransfected cells. Asterisks indicate significance of t-test comparisons to a control GapmeR.

**Figure S2.**
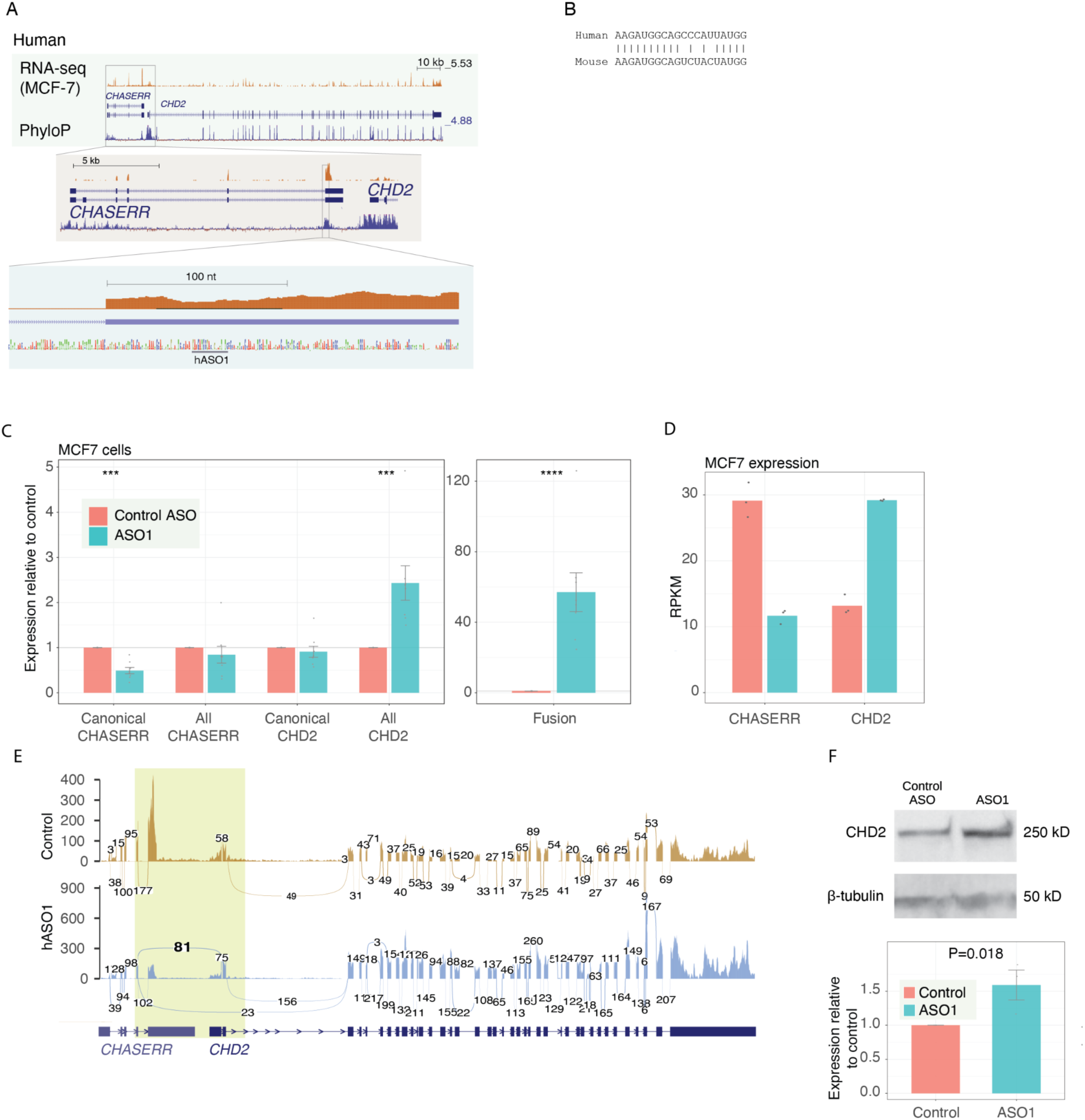
**(A)** As in Fig.1 for the *CHASERR*-*CHD2* locus in the human genome and RNA-seq data from untreated MCF-7 cells. **(B)** Comparison of the sequences of ASO1 and hASO1. **(C)** As in Fig. 1B for human MCF-7 cells. **(E)** As in Fig. 1C for human MCF-7 cells. **(F)** As in Fig. 1E for human MCF-7 cells. **(G)** As in Fig. 1D for human MCF-7 cells.

**Figure S2.**
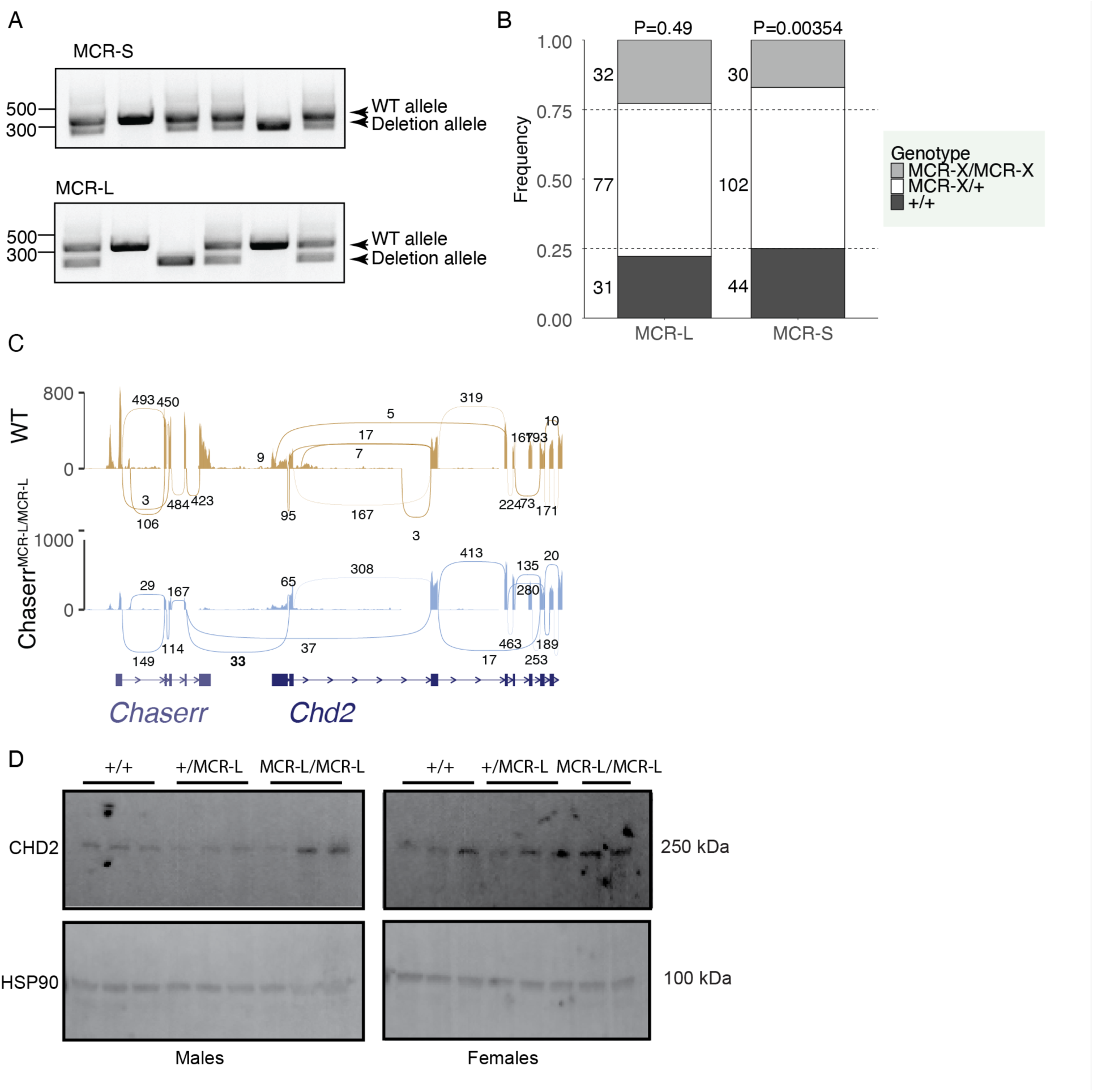
related to Figure 2. Confirmation of fusion formation. **(A)** Sample genotyping PCR gels examining the MCR-S and MCR-L alleles. **(B)** The frequency of different genotypes that were observed following crosses of Chaserr^+/MCR_X^ mice. The number of mice for each genotype is indicated to the left of the bars. P-values represent Chi square deviation from the expected 0.25:0.5:0.25 ratio. **(C)** as in Fig. 2D for the MCR-L allele mice. **(D)** As in Fig. 2F for the MCR-L mice.

**Figure S3.**
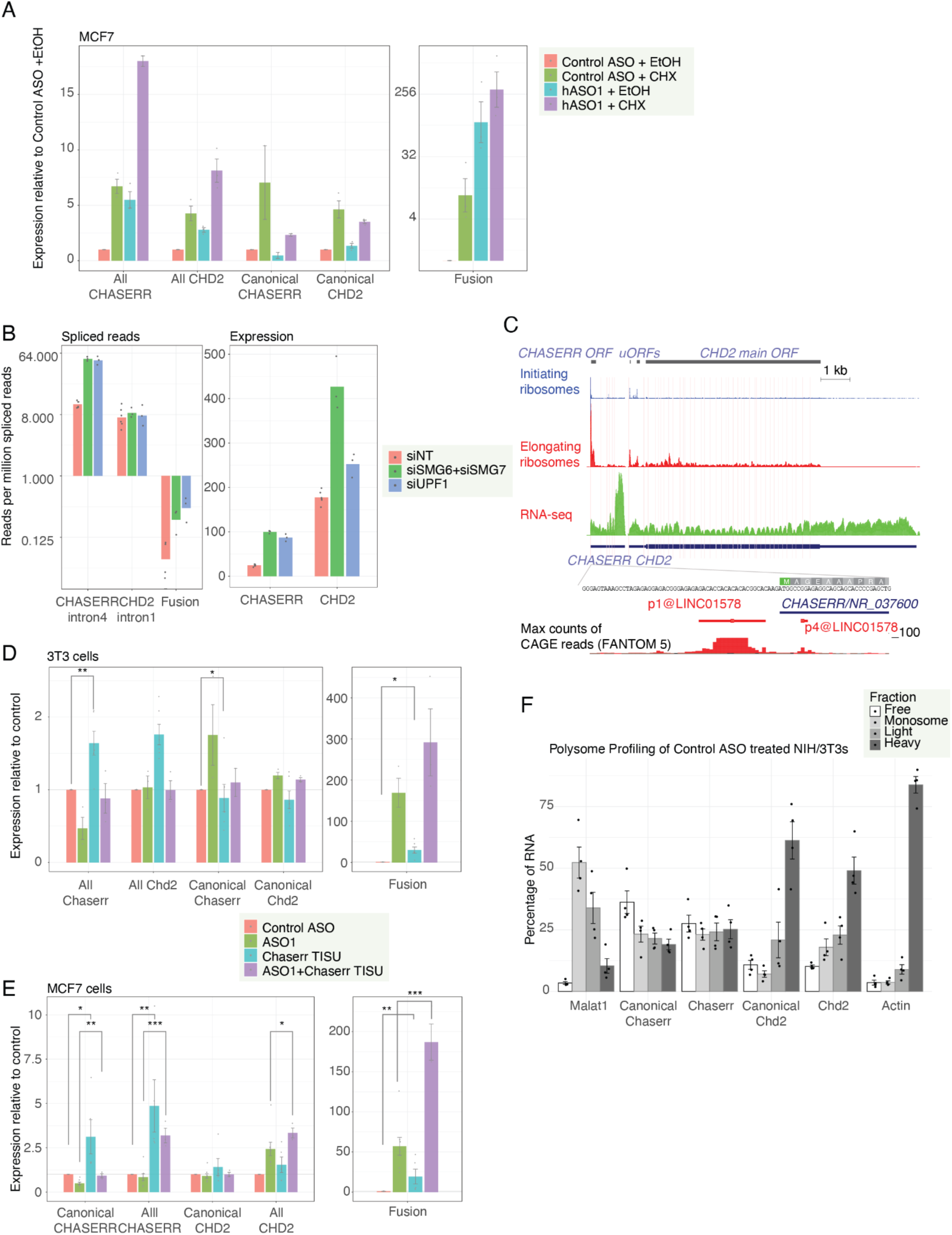
related to Figure 3. NMD sensitivity and translation of the fusion transcript. **(A)** qPCRbased gene expression levels of the indicated transcripts following transfection into MCF-7 cells of the indicated ASOs followed by treatment with either cyclohexamide (CHX) or EtOH vehicle control. Relative expression normalized to β-actin **(B)** Expression levels of the indicated splice junctions (left) or genes (right) from RNA-seq data from^16^ in HeLa cells. **(C)** Alignment of Ribo-seq and RNA-seq reads from aggregated human experiments in GWIPS-VIZ ^13^. The longest ORF in Chaserr, and the two longest uORFs in the Chd2 5’UTR are indicated. Ribo-seq reads are separated into those from regular Ribo-seq libraries sequencing footprints of elongating ribosomes, and ribosomes treated with reagents that arrest them at translation initiation sites. **(D-E)** Changes in expression of the indicated genes following transfection into 3T3 cells (D) or MCF-7 cells (E) of the indicated ASOs. **(F)** As in Fig. 3F for control ASO-treated cells.

**Figure S4.**
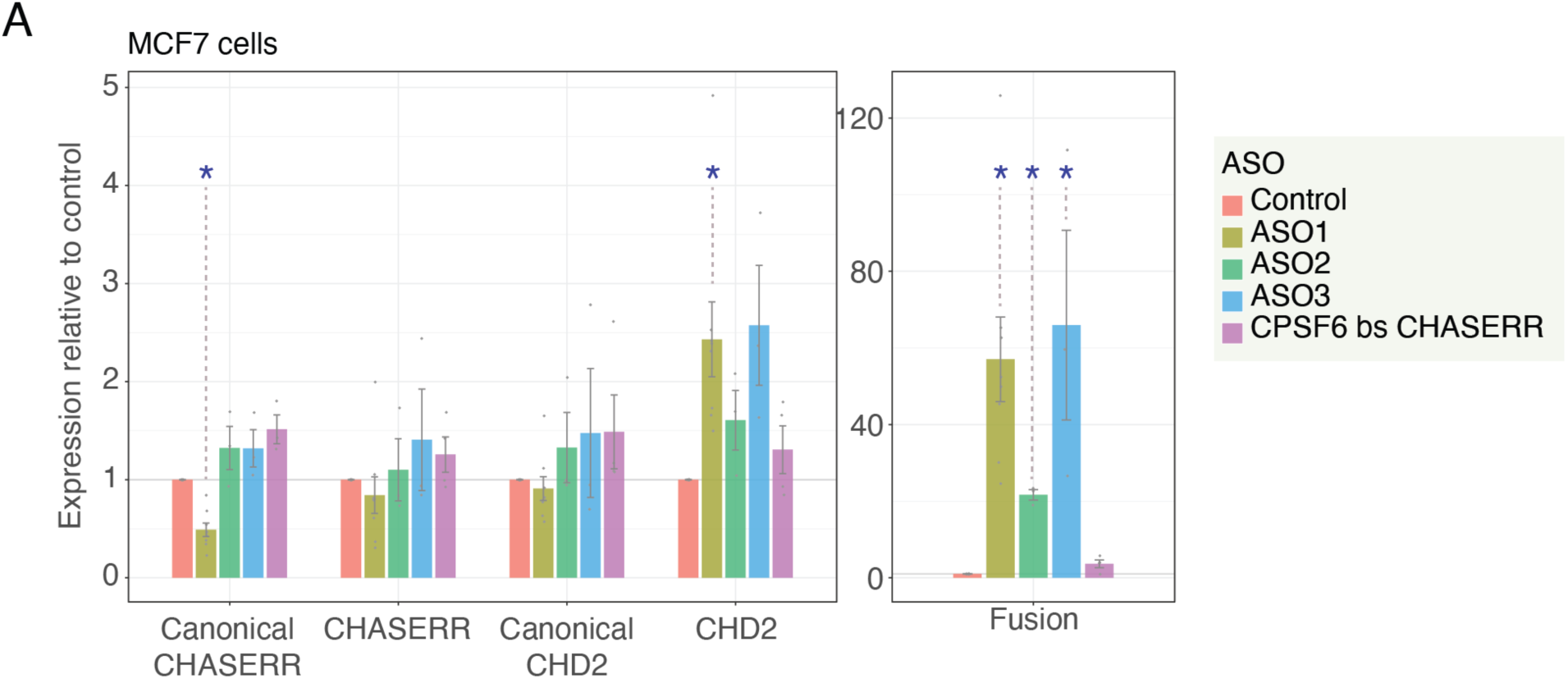
related to Figure 4. **(A)** As in Fig. 4E for ASO treatments in MCF-7 cells.

**Figure S5.**
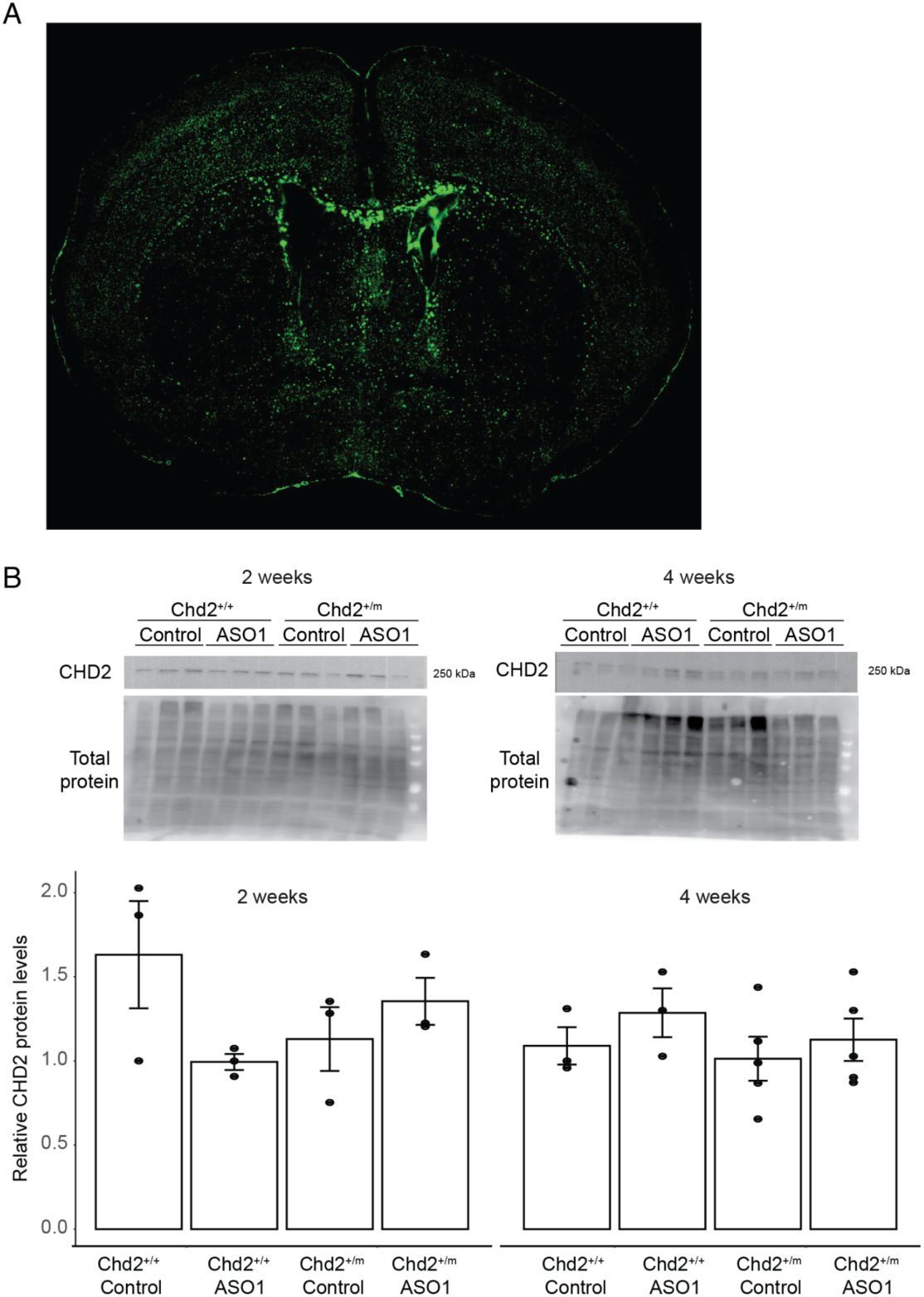
related to Figure 6. ASO injections. **(A)** Distribution of FAM-labelled control ASO in the mouse brain at 2 weeks following injection at P2. **(B)** Western blot (top) and quantification (bottom) of CHD2 protein levels in the whole brain of mice injected at with the indicated ASO at the age of P2 and harvested

**Figure S6.**
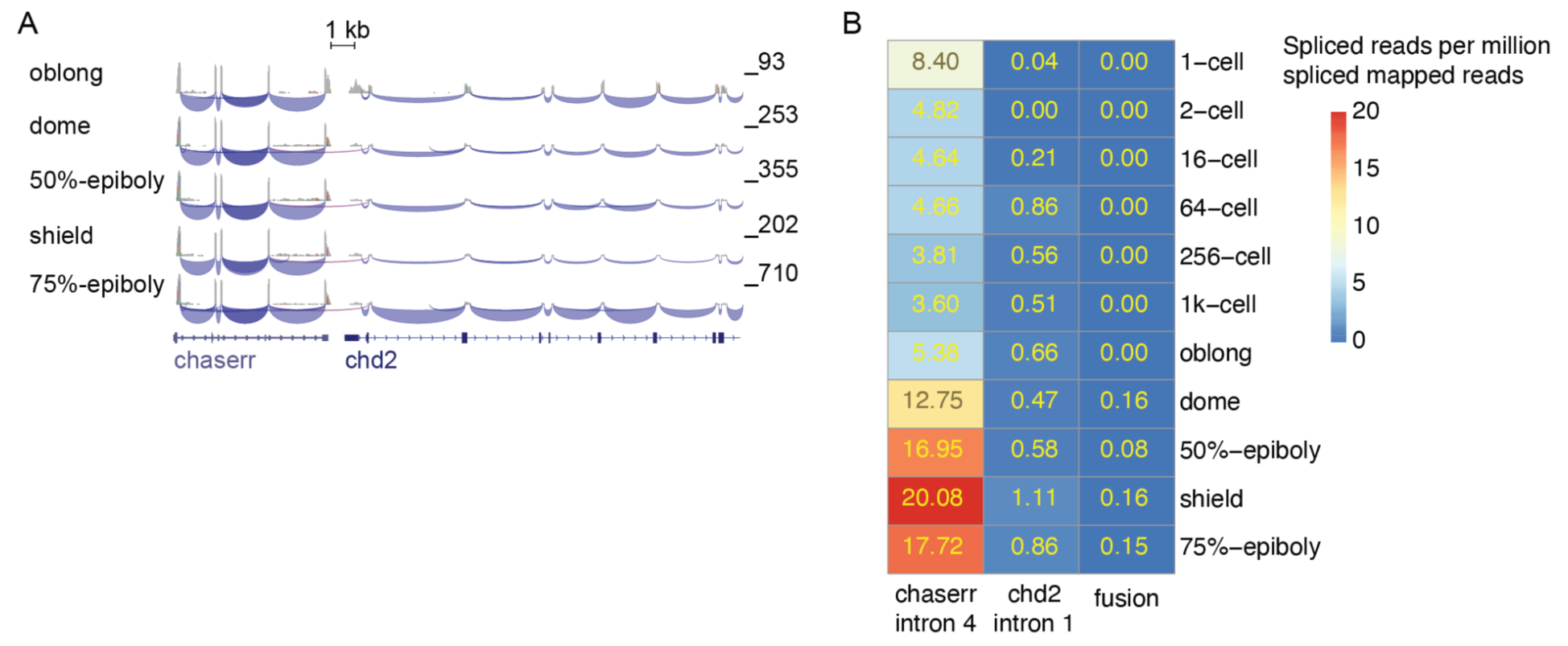
chaserr-chd2 fusion transcript in zebrafish. **(A)** RNA-seq read coverage and spliced reads in the chaserr/chd2 locus (only the first nine exons of chd2 are shown) using data from ^34^. **(B)** Normalized numbers of reads spanning the indicated exon-exon junctions.

